# Aging Of Antiviral CD8^+^ Memory T Cells Fosters Increased Survival, Metabolic Adaptations And Lymphoid Tissue Homing

**DOI:** 10.1101/191932

**Authors:** Bennett Davenport, Jens Eberlein, Verena van der Heide, Kevin Jhun, Tom T. Nguyen, Francisco Victorino, Andrew Trotta, Jerry Chipuk, Zhengzi Yi, Weijia Zhang, Eric T. Clambey, Donald K. Scott, Dirk Homann

**Author notes:** Correspondence: Dirk Homann, MD Diabetes Obesity Metabolism & Immunology Institutes Mount Sinai School of Medicine One Gustave L. Levy Place - Box 1152 New York, NY 10029. BD and JE contributed equally.

## Abstract

Aging of established antiviral T cell memory fosters a series of progressive adaptations that paradoxically improve rather than compromise protective CD8^+^T cell immunity. We now provide evidence that this gradual evolution, the pace of which is contingent on the precise context of the primary response, also impinges on the molecular mechanisms that regulate CD8^+^ memory T cell (CD8^+^T_M_) homeostasis. Over time, CD8^+^T_M_ become more resistant to apoptosis and acquire enhanced cytokine responsiveness without adjusting their homeostatic proliferation rates; concurrent metabolic adaptations promote increased CD8^+^T_M_ quiescence and fitness but also impart the re-acquisition of a partial effector-like metabolic profile; and a gradual redistribution of aging CD8^+^T_M_ from blood and nonlymphoid tissues to lymphatic organs results in CD8^+^T_M_ accumulations in bone marrow, splenic white pulp and particularly lymph nodes. Altogether, these data demonstrate how temporal alterations of fundamental homeostatic determinants converge to render aged CD8^+^T_M_ poised for greater recall responses.

**ABBREVIATIONS:**

AT: adoptive transfer
ATGL: adipose triglyceride lipase
BMP: blood and marginated pool
FA, FAO, FAS, FASN: fatty acid, fatty acid oxidation, fatty acid synthesis, fatty acid synthase
FSC, SSC: forward scatter, side scatter
GMFI: geometric mean of fluorescence intensity
GP, NP: glycoprotein, nucleoprotein
GSEA: gene set enrichment analysis
GSH: glutathione
I^o^, II^o^: primary, secondary
KEGG: Kyoto Encyclopedia of Genes and Genomes
LAL: lysosomal acid lipase (LIPA)
LCMV: lymphocytic choriomeningitis virus
NES: normalized enrichment score
NLTs: nonlymphoid tissues
O, Y: old, young
OxPhos: oxidative phosphorylation
p14 cells: TCRtg CD8^+^T cells specific for the LCMV-GP_33-41_ determinant
RP, WP: red pulp, white pulp (spleen)

T cell subsets

T_E_: effector T cells
T_CM_: central memory T cells (CD62L^hi^)
T_EM_: effector memory T cells (CD62L^lo^)
T_EMRA_: terminally differentiated CD45RA^+^ effector memory T cells (human)
T_M_: memory T cells
T_MP_: memory-phenotype T cells (CD44^hi^)
T_N_: naïve T cells (CD44^lo^)
T_RM_: resident memory T cells (CD69/CD103-enriched)
TCRtg: T cell receptor transgenic
TSLP: Thymic stromal lymphopoietin

## INTRODUCTION

The long-term preservation of antiviral T cell memory is a highly dynamic process that promotes the progressive molecular, phenotypic and functional remodeling of its principal constituents, the populations of specific CD8^+^ memory T cells (CD8^+^T_M_) distributed throughout and often trafficking between various anatomic compartments. We recently demonstrated that this process can culminate, paradoxically, in the acquisition of naïve-like T cell traits, enhanced recall potential and greater protective capacities of aged CD8^+^T_M_ [1]. The notion that aging can improve CD8^+^T cell memory [1-5] stands in apparent contrast to much of the literature documenting numerous and often deleterious consequences of T cell aging [6-9]. A more focused review [10], however, indicates that eventual “immunosenescence” is not necessarily a fate shared by all T cell subsets and CD8^+^T_M_ generated earlier in life to an acute, non-persisting pathogen challenges can be maintained over time without accruing obvious functional defects [9, 11, 12]. In fact, a coexistence of age-associated alterations that either impair or improve CD8^+^T cell immunity is illustrated in aged mice that exhibit a diminished capacity for generation of primary (I^o^) antiviral CD8^+^ effector T cell (CD8^+^T_E_) responses yet readily support the greater secondary (II^o^) expansion of old as compared to young CD8^+^T_M_ specific for the same viral determinants [1].

To elucidate the foundations and consequences of successful CD8^+^T_M_ aging in greater detail, we previously generated a set of integrated data sets that collectively trace the evolving molecular, phenotypic and functional properties of aging virus-specific CD8^+^T_M_ [1]. We further organized the patterns of gradual CD8^+^T_M_ remodeling in a conceptual framework designated the “rebound model” of progressive CD8^+^T_M_ “de-differentiation” that postulates an inverse relationship between the extent of I^o^ CD8^+^T_E_ differentiation and the pace with which aging CD8^+^T_M_ populations, over a period of ∼2 years, acquire a broad spectrum of distinctive and increasingly homogenous traits. Specifically, aging of CD8^+^T_M_ populations modulates the expression of at least ∼80 cell surface receptors/ligands, produces a more diversified functional repertoire, and eventually endows old CD8^+^T_M_ in a T cell-intrinsic fashion with an improved capacity for the generation of protective II^o^ responses [1]. In the present report, we have delineated the impact of aging on the cardinal components of CD8^+^T_M_ homeostasis (survival, homeostatic proliferation, metabolism, tissue residence/trafficking) and our findings demonstrate that the cumulative homeostatic adaptations converge to establish a spatio-functional foundation for improved recall responses of aged CD8^+^T_M_.

## RESULTS AND DISCUSSION

### Temporal regulation of survival-& apoptosis-related gene and protein expression by aging CD8^+^T_M_

Challenge of mice with the natural murine pathogen lymphocytic choriomeningitis virus (LCMV) induces a potent antiviral CD8^+^T_E_ response that rapidly controls the infection and permits the subsequent development of specific CD8^+^T_M_ that are maintained for life in the absence of residual viral antigens [13, 14]. In B6 mice, the principal LCMV-specific CD8^+^T cell populations target the nucleo- and glycoprotein determinants NP_396-404_ and GP_33-41_; in addition, naïve TCRtg p14 cells specific for LCMV-GP_33-41_ and transferred into congenic B6 mice can be used to construct “p14 chimeras” for facilitated interrogation of a clonotypic CD8^+^T_M_ population (p14 T_M_). A combination of p14 chimera and B6 systems provided the experimental foundation for our comprehensive delineation of aging antiviral CD8^+^T_M_ properties [1], and drawing on these resources, we have now revisited the foundations of long-term CD8^+^T_M_ survival [15] by conducting modified gene set enrichment analyses (GSEAs) that specifically leverage the temporal aspect of our p14 T_M_ data sets (see Methods). Here, of 132 gene sets comprising 10,945 genes and exhibiting age-associated modulations, ∼20% (26/132) were enriched and ∼80% (106/132) were depleted in old p14 T_M_, the latter group including the KEGG apoptosis pathway (***Fig.1A***). Within this module, 14 genes belonged to the Bcl-2, BIRC (baculoviral IAP [inhibitors of apoptosis proteins] repeat-containing) or caspase gene families and their combined temporal regulation pointed towards reduced apoptosis susceptibility of aged p14 T_M_ (***Fig.1A***).

**Figure 1.**
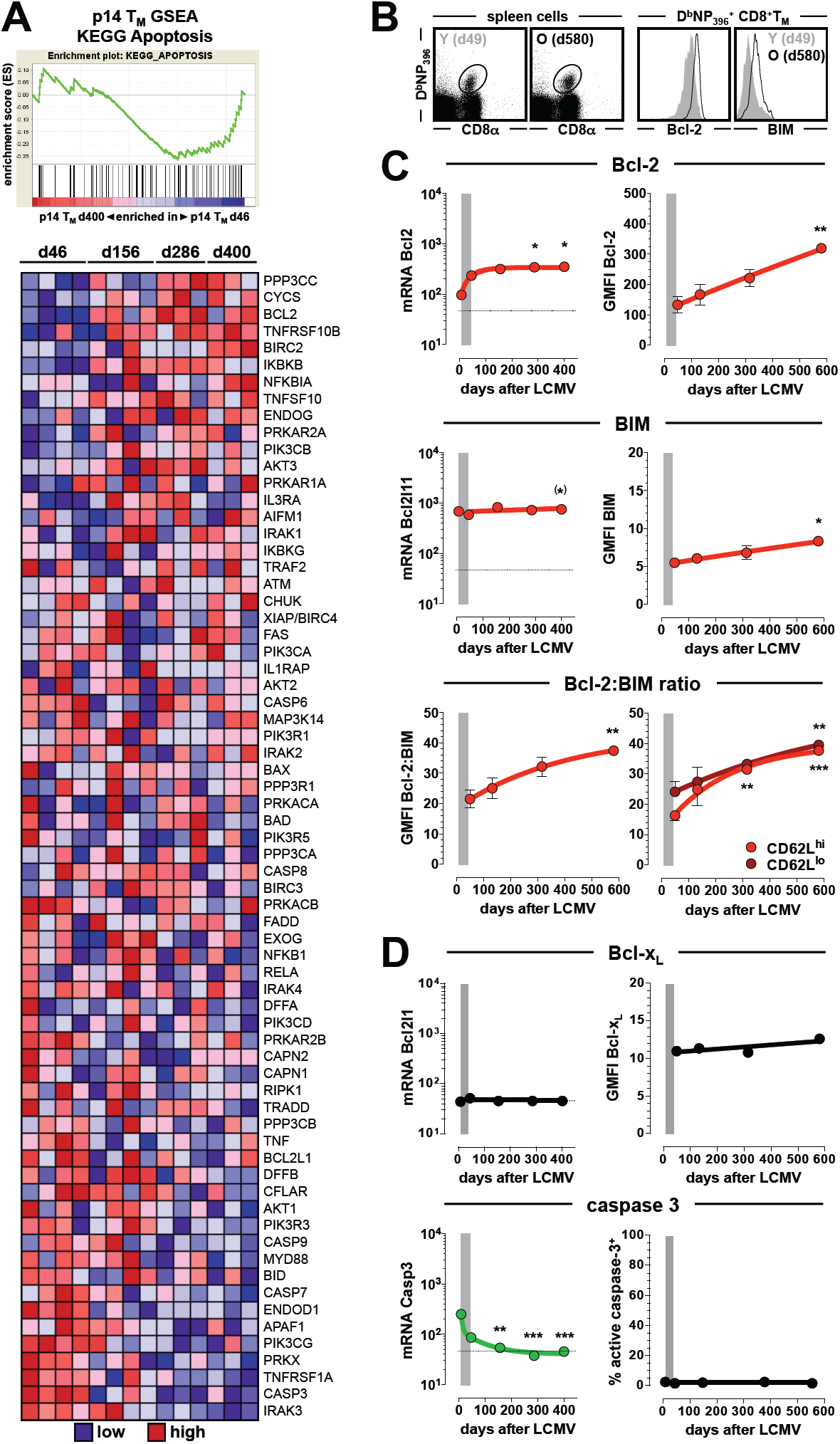
Temporal regulation of major survival-associated components by aging CD8^+^T_M_. **A.,** GSEAs were performed with p14 T_M_ data sets (d46-d400 after virus challenge) as described in Methods and demonstrate a relative depletion of genes within the KEGG apoptosis module for aged p14 T_M_ (normalized enrichment score [NES]: -1.04); the corresponding heat map displays relative expression levels of GSEA-ranked genes within that module. **B.,** staining/gating strategy and representative Bcl-2 and BIM expression data in young and old CD8^+^T_M_. **C. & D.,** progressive modulation of survival/apoptosis-related mRNA (p14T_E/M_) and protein (D^b^NP_396_^+^ CD8^+^T_E/M_) expression levels; Bcl-2:BIM ratios were calculated by division of respective GMFI (geometric mean of fluorescence intensity) values and are shown for both total D^b^NP_396_^+^ CD8^+^T_M_ and subsets stratified according to CD62L expression. The vertical gray bars indicate the transition period from CD8^+^T_E_ stage (d8) to early T_M_ stage (d42), and significant differences emerging over the course of the memory phase (comparing young and older specific CD8^+^T_M_ populations by one-way ANOVA with Dunnett’s multiple comparisons test) are highlighted in red (up-regulation) or green (down-regulation); the parenthetical asterisk in the *Bcl2l11* graph indicates significance between d46 and d400 p14T_M_ as calculated by Student’s t-test (n≥3 individual mice per time point and experiment).

Members of the Bcl-2 family have long been implicated in lymphocyte survival and death, and the balanced expression of anti-apoptotic Bcl-2 and pro-apoptotic BIM controls survival of naïve and, to a somewhat lesser extent, memory phenotype CD8^+^T cells (CD8^+^T_N_ and CD8^+^T_MP_, respectively) [16]. Our interrogation of individual Bcl-2 family members revealed predominantly stable expression by aging p14 T_M_ with two notable exceptions, the modestly rising levels of *Bcl2* and *Bcl2l11* (***Fig.S1A-C***). Importantly, the transcriptional changes were accompanied by a substantial increase of Bcl-2 protein content in aging CD8^+^T_M_ and a slight, though significant, enhancement of BIM such that the resulting Bcl-2:BIM expression ratio steadily increased over time (***Fig.1B/C***). Given a progressive enrichment for the CD62L^hi^ phenotype among aging antiviral CD8^+^T_M_ [1, 17], our findings are also in agreement with the reported elevation of both Bcl-2 and BIM in antiviral CD8^+^T_CM_ as compared to T_EM_ populations [18]. We emphasize, however, that these expression differences themselves are subject to an extended temporal modulation since the continuous rise of the Bcl-2:BIM ratios occurred in aging CD8^+^T_CM_ and T_EM_ subsets alike (***Fig.1C***). We also note the persistence of relatively stable Bcl-x_L_ levels (***Fig.1D***); a gradual increase of several BIRC family genes that may contribute to an enhanced survival advantage for aged CD8^+^T_M_ [19, 20] (***Fig.S1D***); and the pronounced decline of *Casp3* mRNA without evidence for caspase-3 activation [21] throughout long-term T cell memory (***Figs.1D & S1E***). Altogether, the kinetics of gene and protein expression therefore indicate that aging CD8^+^T_M_ may be endowed with increasing overall fitness.

### Enhanced apoptosis resistance of aging CD8^+^T_M_

When assessed directly *ex vivo*, the viability of CD8^+^T_M_ was not affected by age (***Fig.2A***), but an *in vitro* culture in the absence of added growth/survival factors (“withdrawal apoptosis”) documented a gradual decline of CD8^+^T_M_ death as a function of age (***Fig.2B***). Increased apoptosis resistance has been associated with aging and cellular senescence [22] but the CD8^+^T_M_ under investigation here lacked phenotypic and functional features of incapacitation [1], including the hallmark of murine T cell senescence, increased P-glycoprotein activity [23]. Nonetheless, a distinct survival advantage of “non-senescent” old CD8^+^T_MP_ was previously observed under conditions of “withdrawal apoptosis” and attributed, despite an exacerbated decline of mitochondrial membrane potentials (ΔΨm), to reduced production of reactive oxygen species (ROS), elevated intracellular thiol levels (largely representing the abundance of reduced glutathione/GSH), and increased expression of phase II antioxidant enzymes that combine to protect aged CD8^+^T_MP_ against oxidative stress, mitochondrial dysfunction and death [24, 25]. In our model system, aging virus-specific CD8^+^T_M_ also exhibited a modest decline of *ex vivo* ROS production (***Fig.2C***) and a more striking loss of ΔΨm after *in vitro* culture (***Fig.2D***). Yet despite an enrichment of genes within the GSH metabolism pathway (***Fig.2E***) that may collectively provide a metabolic advantage [26] for recall responses, we observed only a marginal rise of intracellular thiol levels in aging CD8^+^T_M_ (***Fig.2F***), and regardless of a 1.9-fold increase of *Nfe2l2* mRNA [1] (the major TF in control of phase II enzyme regulation), no evidence for elevated induction of the respective genes could be obtained (not shown). Instead, we found a pronounced augmentation of cell surface thiol levels by aging CD8^+^T_M_ that was likely the result of changing microenvironments in older mice as demonstrated by their significantly increased serum thiol levels (***Fig.2G***); this conclusion is also consistent with the notion that the immediate microenvironment rather than intracellular GSH levels preferentially determines the redox status of cell surface molecules [27].

**Figure 2.**
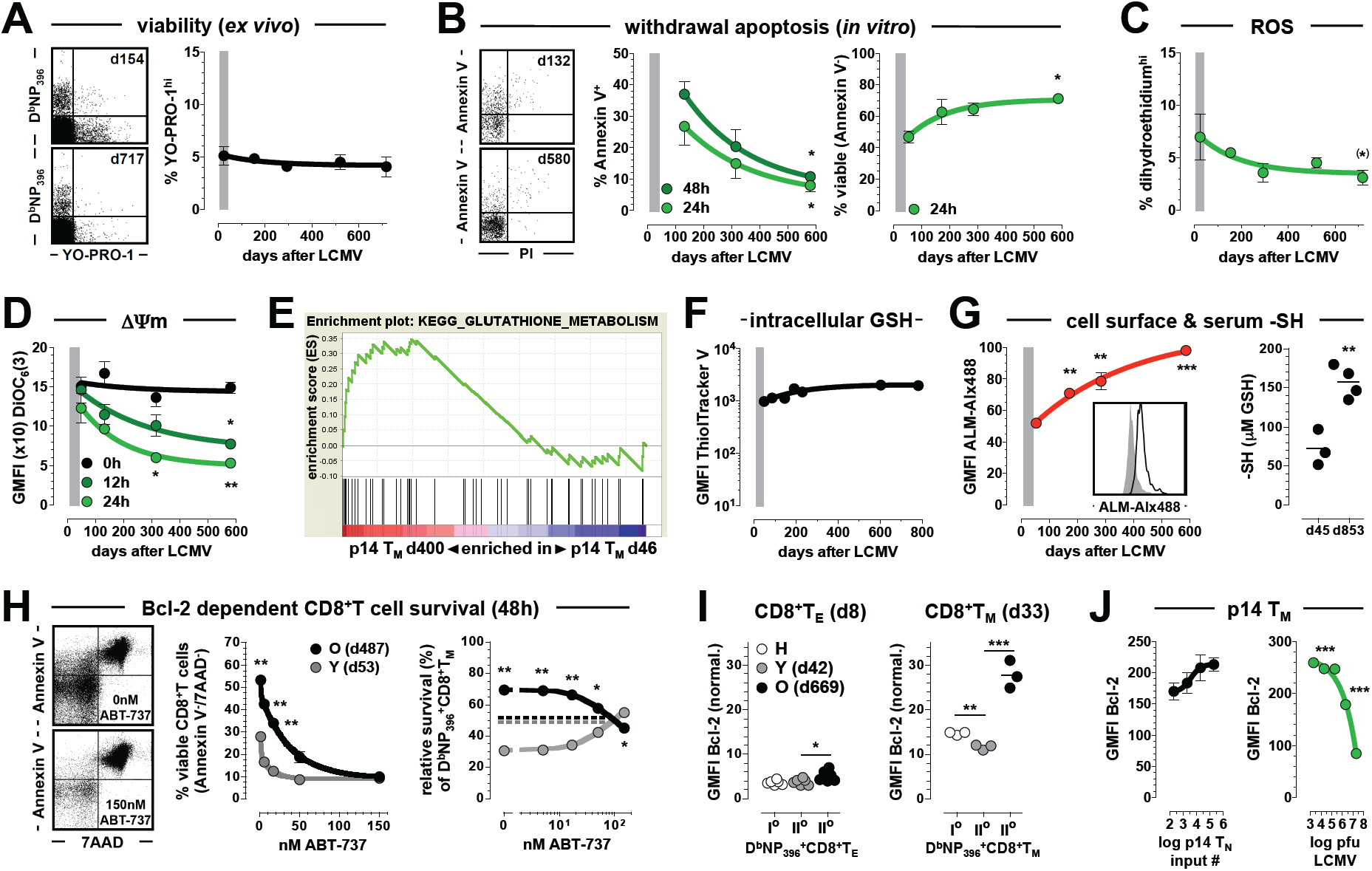
Life & death of aging CD8^+^T_M_. **A.,** viability of blood-borne D^b^NP_396_^+^ CD8^+^T_M_ as assessed directly *ex vivo* (dot plots gated on CD8^+^T cells). **B.,** survival of splenic NP_396_-specific CD8^+^T_M_ was determined after 24-48h *in vitro* culture in the absence of added survival/growth factors (“withdrawal apoptosis”, dot plots gated on D^b^NP_396_^+^ CD8^+^T_M_); data from 2 separate experiments display apoptosis/death (middle) or survival (right) of D^b^NP_396_^+^ CD8^+^T_M_ as a function of age. **C.,** reactive oxygen species (ROS) production capacity of blood-borne D^b^NP_396_^+^ CD8^+^T_M_. **D.,** mitochondrial membrane potential (⊗Ψm) of D^b^NP_396_^+^ CD8^+^T_M_ was measured as a function of time after LCMV challenge and duration of *in vitro* culture (0-24h). **E.,** GSEA analysis of glutathione (GSH) metabolism (normalized enrichment score [NES]: 1.28). **F.,** intracellular GSH levels of aging blood-borne D^b^NP_396_^+^ CD8^+^T_M_. **G.,** modulation of cell surface thiol levels by aging D^b^NP_396_^+^ CD8^+^T_M_ as determined by maleimide-Alx488 staining (the insert compares young [gray: d43] and old [black: d575] D^b^NP_396_^+^ CD8^+^T_M_); plasma thiol groups were quantified in young and old LCMV-immune B6 mice as indicated using 5,5’-dithiobis(2-nitrobenzoic acid), and data are expressed in relation to a GSH standard. **H.,** CD8^+^T cells enriched from young and old congenic mice were mixed 1:1 at the level of D^b^NP_396_^+^ CD8^+^T_M_ and cultured for 48h in the absence or presence of the Bcl-2 antagonist ABT-737. Left: dot plots gated on total CD8^+^T cells; middle: viability of young *vs.* old CD8^+^T cells as a function of ABT-737 concentration; right: survival of D^b^NP_396_^+^ CD8^+^T_M_ is displayed as the relative preponderance of young *vs.* old populations after 48h of culture (the dotted line indicates the original input ratio of Y:O = 49:51%). **I.,** Bcl-2 expression levels of blood-borne I^o^ (H: host) and II^o^ (Y *vs.* O) D^b^NP_396_^+^ CD8^+^T_E/M_ generated in the same animals and analyzed on d8 (left) and d33 (right) after mixed AT/re-challenge. **J.,** Bcl-2 expression by p14 T_M_ (d44-49) as a function of original p14 T_N_ input number (left) or LCMV challenge dosage (right); n≥3 individual mice per time point in 2-4 independent experiments.

### Life & death of aging CD8^+^T_M_: improved survival through increased Bcl-2:BIM expression ratios

Collectively, the above observations suggest that T cell-intrinsic mechanisms, in particular the rising Bcl-2:BIM expression ratio, may confer a survival advantage to aging CD8^+^T_M_. To directly evaluate this possibility we employed a co-culture system to monitor survival of congenic old and young CD8^+^T_M_ in the same *in vitro* environment. Addition of the Bcl-2 inhibitor ABT-737 [28] to cultures precipitated CD8^+^T cell death in a dose-dependent fashion and, at a saturating concentration of 150nM, reduced total CD8^+^T cell survival to ∼10% (***Fig.2H***). The relative survival advantage of old *vs.* young D^b^NP_396_^+^ CD8^+^T_M_, however, was maintained at lower ABT-737 dosages and only disappeared at ∼100nM providing direct evidence for the exquisite dependence of CD8^+^T_M_ survival on Bcl-2 and its role in promoting enhanced apoptosis resistance of aged CD8^+^T_M_ populations (***Fig.2H***). Although ABT-737 binds to Bcl-x_L_ and Bcl-w in addition to Bcl-2 [28], the low-level expression of corresponding mRNA species and, in the case of Bcl-x_L_ also protein (***Figs.S1A & 1C***), supported the notion of Bcl-2 as the major ABT-737 target in CD8^+^T_M_. Further evidence for the elevated Bcl-2:BIM ratio as a determinant for enhanced survival of aged CD8^+^T_M_ came from a reversal of survival advantages at saturating ABT-737 concentrations (150nM, ***Fig.2H***): although very few cells remained alive under conditions of complete Bcl-2 blockade, the slightly better survival of residual young CD8^+^T_M_ may be explained by their comparatively lower BIM expression since death of Bcl-2-deficient T cells was shown earlier to decline as a function of BIM gene dosage (*Bcl2l11*^+/+^ > *Bcl2l11*^+/-^ > *Bcl2l11*^-/-^) [16].

In the context of an acute response, both I^o^ and II^o^ CD8^+^T_E_ downregulate Bcl-2 expression [29], and control of CD8^+^T_E_ subset survival is thought to switch to other factors, perhaps including the BIRC family member survivin/Birc5 [30] (***Fig.S1D***). Work with a Bcl-2 reporter mouse, however, indicates that even at the peak of a pathogen-specific immune response, CD8^+^T_E_ populations are characterized by a spread of Bcl-2 expression levels that permits the distinction of CD8^+^T_E_ subsets with differential memory potential [31]. In line with these observations, we found that II^o^ CD8^+^T_E_ derived from aged CD8^+^T_M_ exhibited a slight yet significant elevation of Bcl-2 as compared to I^o^ CD8^+^T_E_ or II^o^ CD8^+^T_E_ generated from young CD8^+^T_M_ (***Fig.2I***). Coupled with the former cells’ improved survival during the ensuing contraction phase [1], our results therefore hinted at a direct role for Bcl-2 in promoting a more effective establishment of II^o^ CD8^+^T cell memory. Indeed, while young II^o^ CD8^+^T_M_ featured reduced Bcl-2 contents compared to I^o^ CD8^+^T_M_ as reported previously [29], old II^o^ CD8^+^T_M_ present within the same hosts exhibited substantially higher Bcl-2 levels (***Fig.2I***). Thus, the largely Bcl-2-dependent survival advantage of old over young I^o^ CD8^+^T_M_ was re-established in the course of II^o^ memory formation.

Overall, the dynamic regulation of Bcl-2 re-expression in the memory phase (***Fig.1B/C***) followed a pattern similar to that of multiple other phenotypic/functional CD8^+^T_M_ properties subject to age-associated expression modulation [1]. Since the precise pace of these changes could be experimentally accelerated or delayed as a function of initial CD8^+^T_N_ precursor frequency or infection dosage [1], we surmised that Bcl-2 expression by CD8^+^T_M_ could be controlled in a comparable fashion. Here, we constructed p14 chimeras with titered numbers of p14 T_N_ (2×10^2^ – 2×10^5^) and challenged the mice with a standard dose of LCMV (2×10^5^ pfu), or generated p14 chimeras with a fixed p14 T_N_ number (1×10^4^) and infection with graded dosages of LCMV (2×10^3^ – 2×10^7^ pfu). Measuring Bcl-2 expression by p14 T_M_ 6-7 weeks later, we found that an increase of p14 T_N_ input numbers enhanced, while an escalation of the virus challenge dose reduced respective Bcl-2 levels in p14 T_M_ (***Fig.2J***). In summary, our results demonstrate that aging CD8^+^T_M_ become more resistant to apoptosis, that their improved survival and that of their II^o^ progeny is principally controlled through increased Bcl-2 expression, and that the specific conditions of CD8^+^T_E_ generation determine the pace of progressive Bcl-2 upregulation by CD8^+^T_M_.

### Cytokine receptor expression, signaling & homeostatic proliferation of aging CD8^+^T_M_

In direct relation to their longevity, regulation of CD8^+^T_M_ fates under steady-state conditions also involves homeostatic proliferation, the slow and stochastic division of “resting” CD8^+^T_M_ governed by the cytokines IL-7 and IL-15 [15, 32]. In extension of our previous report [1], we now demonstrate that a progressive upregulation of the respective cytokine receptors (CD127 and CD122) by aging CD8^+^T_M_ also pertains to the PBMC compartment and to differential CD8^+^T_M_ specificities (***Fig.3A/B***) suggesting that their homeostatic proliferation rates may be adjusted accordingly. To determine if enhanced cytokine receptor expression indeed conveyed greater responsiveness, we assessed the extent of IL-7/IL-15-induced STAT5 phosphorylation in young and old p14 chimeras. Here, aged p14 T_M_ not only exhibited greater reactivity, but at limiting concentrations IL-7 clearly proved to be a more effective activator of STAT5 than IL-15 (***Fig.3C***). These findings extend the notion of superior IL-7 potency in the context of initial CD8^+^T_M_ formation [33] to the long-term maintenance of CD8^+^T_M_, and complement a recent observation about enhanced IL-15 reactivity of “late” p14 T_M_ or T_CM_ (>8 months after infection) as compared to “early” p14 T_M_/T_CM_ (d30-45) [5]. We further note that the thymic stromal lymphopoietin receptor (TSLPR) is apparently the only CD8^+^T_M_-expressed cytokine receptor subject to a gradual downmodulation over time [1], a pattern that could contribute to the amplified IL-7 reactivity of aged CD8^+^T_M_ as it may permit enhanced complex formation of CD127 with CD132 rather than TSLPR [34].

**Figure 3.**
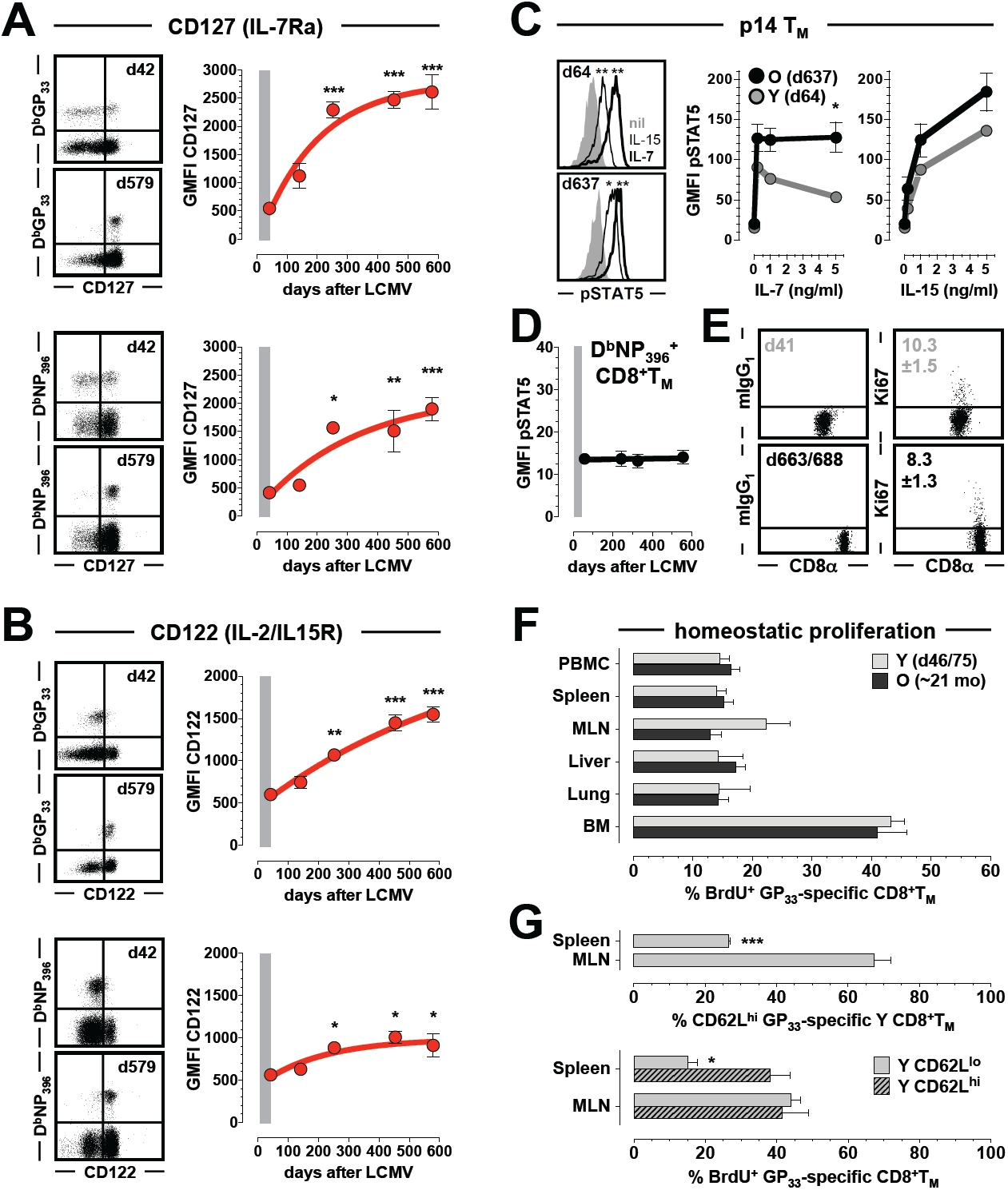
CD127/CD122 expression, signaling and homeostatic proliferation of aging CD8^+^T_M_. **A.,** cohorts of young adult B6 mice were challenged with LCMV in a staggered fashion and contemporaneous analyses of aging CD8^+^T_M_ populations were conducted with peripheral blood. Dot plots are gated on CD8^+^T cells and display CD127/IL-7Ra expression by young and old D^b^GP_33_^+^ (top) and D^b^NP_396_^+^ (bottom) CD8^+^T_M_; note that data for D^b^GP_33_^+^ and D^b^NP_396_^+^ CD8^+^T_M_ were generated with different flow cytometers such that GMFI values between these populations cannot be directly compared (n=4 mice/time point). **B**., temporal regulation of CD122/IL-2Rb (also part of the IL15R complex) expression by blood-borne D^b^GP_33_^+^ and D^b^NP_396_^+^ CD8^+^T_M_; data organization as in panel A. **C.,** IL-7 and IL-15 responsiveness of young and old p14 T_M_ as determined by STAT5 phosphorylation (15min *in vitro* cytokine exposure); histograms are gated on p14 T_M_ (gray: no cytokine, thin black tracing: IL-15 [0.2ng/ml], thick black line: IL-7 [0.2ng/ml]). Note that maximal respective STAT5 phosphorylation required 0.2ng/ml rIL-7 but ∼10ng/ml rIL-15 (not shown). **D.,** *ex vivo* pSTAT5 levels of aging CD8^+^T_M_. **E.,** Ki67 expression by young and old blood-borne D^b^NP_396_^+^ CD8^+^T_M_ (values indicate average percentage of Ki67^+^ cells [n=5-9 mice; p=ns]). **F.,** homeostatic proliferation of GP_33_-specific CD8^+^T_M_ in different tissues of young and old LCMV-immune B6 mice was assessed with a 7-day *in vivo* BrdU pulse (combined data from 2 independent experiments). **G.,** frequency (top) and homeostatic proliferation (bottom, 7-day BrdU pulse) of CD62L^hi^ and CD62L^lo^ GP_33_-specific CD8^+^T_M_ subsets in spleen and MLN of young LCMV-immune B6 mice (n≥3 individual mice per time point and experiment). Statistical differences were calculated using one-way ANOVA with Dunnett’s multiple comparisons test (panels A/B), or Student’s t-test (panels C-G).

While the above findings correlate increased CD127/CD122 expression with CD8^+^T_M_ reactivity to IL-7/IL-15, we also noted a certain extent of constitutive STAT5 phosphorylation among p14 T_M_ analyzed directly *ex vivo*, similar to the basal STAT5 phosphorylation observed in human CD8^+^T cell subsets of undefined specificity [35]. Additional control experiments confirmed this conclusion (***Fig.S2A***) but unexpectedly, the levels of constitutive STAT5 phosphorylation remained unaltered in aging antiviral CD8^+^T_M_ populations (***Fig.3D***). Since the level of active STAT5 appears to control homeostatic proliferation rates [36], stable pSTAT5 expression by endogenously generated CD8^+^T_M_ therefore suggested that their homeostatic proliferation rates, despite enhanced sensitivity to IL-7/IL-15, might not be accelerated. This prediction was reinforced by our longitudinal p14 T_M_ GSEAs that demonstrated a negative (though not significant) enrichment of cell cycle-associated genes and thus also argued against an accelerated CD8^+^T_M_ turnover (***Fig.S2B***). Indeed, as assessed by *ex vivo* Ki67 expression, homeostatic proliferation of blood-borne LCMV-specific CD8^+^T_M_ was unaffected by age (***Fig.3E***), a contention corroborated through the comparable *in vivo* BrdU incorporation by young and old CD8^+^T_M_ in various lymphatic and nonlymphoid tissues (NLTs) (***Fig.3F***). Thus, in contrast to murine CD8^+^T_MP_ of undefined specificity [37], homeostatic proliferation rates of virus-specific CD8^+^T_M_ were largely independent of age but remained susceptible to modulation by tissue-specific microenvironments as shown for young CD8^+^T_M_ [38].

Finally, it is important to note that homeostatic proliferation rates are not simply an intrinsic property of phenotypically defined CD8^+^T_M_ subsets. For example, the CD62L^hi^ CD8^+^T_CM_ population, previously reported to exhibit higher homeostatic proliferation rates than CD8^+^T_EM_ [17, 39], accumulates in the spleen over time [1, 17] without causing an overall acceleration of homeostatic turnover (***Fig.3E/F***). And although we confirmed the differential homeostatic proliferation rates of splenic CD8^+^T_CM_ *vs.* T_EM_ in young LCMV-immune mice, we found no differences in other tissues such as LNs (***Fig.3G***). The absence of a simple correlation between CD8^+^T_M_ subsets, rates of homeostatic proliferation, cytokine receptor (CD127/CD122) and even corresponding tissue-specific cytokine (*Il7/Il15*) expression levels (***Fig.S2C***) constitutes an important caveat that needs to inform further investigations into the homeostasis of CD8^+^T_M_ populations.

### Metabolic adaptations of aging CD8^+^T_M._

Initial CD8^+^T_E_ differentiation and CD8^+^T_M_ generation are both controlled and accompanied by varied metabolic adaptations. Activation of “quiescent” naïve CD8^+^T_N_ engages a “metabolic switch” that endows emerging CD8^+^T_E_ with high rates of aerobic glycolysis and glutaminolysis to support an anabolic metabolism; the subsequent development of CD8^+^T cell memory is characterized by a gradual return to metabolic quiescence and a preferential reliance on fatty acid oxidation (FAO) and oxidative phosphorylation (OxPhos) to meet the changing energy demands [40]. The extent to which established CD8^+^T_M_ populations may further adapt their metabolism over time, however, remains little explored [5]. We previously reported that aging CD8^+^T_M_ exhibit a subtle yet significant increase of cellular size and “granularity/complexity” (determined by forward [FSC] and side scatter [SSC] properties, respectively) [1], a process most likely controlled by mTOR activity [41, 42]. Indeed, we now find that basal mTOR protein (though not mRNA) expression by antiviral CD8^+^T_M_ increased with age as did message for ribosomal protein S6 (*Rps6*, a downstream target of the mTORC1 complex involved in the regulation of cell size, proliferation and glucose homeostasis) and, importantly, the degree of Rps6 protein phosphorylation (***Fig.4A/B***). Although the convergence of elevated mTORC1 activity, cell size and recall capacity of aged CD8^+^T_M_ [1] is consistent, these adjustments would appear to run counter to the shift towards reduced glycolysis and increased OxPhos as observed for the earlier transition from CD8^+^T_E_ to young T_M_ stage [40]. Interestingly, however, most recent work indicates that enforcement of sustained glycolysis and suppression of OxPhos does not compromise but rather may accelerate CD8^+^T_M_ formation [43]. Therefore, to assess the extended evolution of metabolic CD8^+^T_M_ profiles, we reviewed our temporal GSEAs and found that ∼25% of all pathways up- or downregulated by p14 T_M_ over time could in fact be assigned to the broad KEGG category of “metabolism”. Here, a collective depletion of carbohydrate, energy, lipid, amino acid and glycan pathways in aging p14 T_M_ suggested a continued trend towards metabolic quiescence yet the gene sets comprising glycolysis, nucleotide and glutathione metabolism were simultaneously enriched (***Fig.4C*** and not shown). In the absence of significant differences for the majority of these temporally regulated pathways (***Fig.4C***), the age-associated alterations of CD8^+^T_M_ metabolism are therefore expected to be subtle but nevertheless should be reflected in a distinct modulation of glucose and fatty acid utilization.

**Figure 4.**
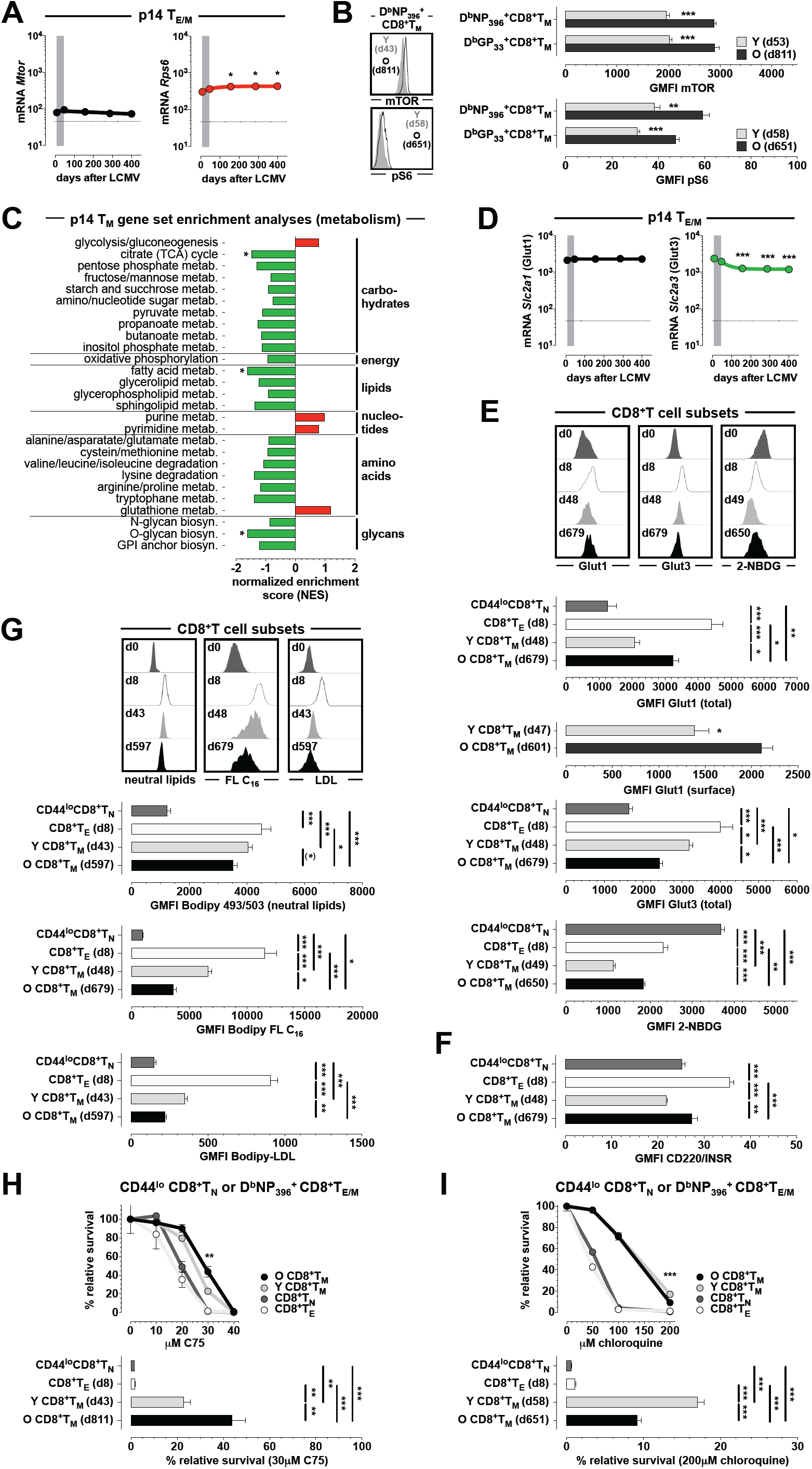
Metabolic adaptations of aging CD8^+^T_M_. **A.,** temporal regulation of *Mtor* and *Rps6* expression by aging p14 T_M_. **B.,** expression of mTOR and phosphorylated Rps6 (pS6) by young and old LCMV-specific CD8^+^T_M_. **C.,** GSEAs were conducted with previously generated data sets on aging p14 T_M_ as detailed in Methods, and the panel summarizes the temporally regulated gene sets progressively enriched or depleted within the KEGG metabolism module (statistical significance in only three pathways is indicated by asterisks). **D.,** temporal regulation of *Slc2a1* and *Slc2a3* expression by aging p14 T_M_. **E.,** expression levels of total Glut1 (intracellular stain), surface Glut1 (Glut1.RBD.GFP stain) or total Glut3 were determined for CD44^lo^CD8^+^T_N_ (d0), D^b^NP_396_^+^ CD8^+^T_E_ (d8) as well as indicated young and old D^b^NP_396_^+^ CD8^+^T_M_ in multiple contemporaneous experiments conducted with splenic or blood-borne CD8^+^T cell populations (histograms are gated on indicated “live” [zombie^-^] CD8^+^T cell subsets). Bottom panel: glucose uptake by indicated CD8^+^T cell populations was quantified using the fluorescently-labeled deoxyglucose analog 2-NBDG. **F.,** expression of insulin receptor (CD220) by indicated CD8^+^T cell populations. **G.,** neutral lipid content as well as long-chain FA and LDL uptake by indicated CD8^+^T cell populations was quantified using Bodipy 493/503, Bodipy FL C16 or Bodipy-LDL staining, respectively (overall experimental design and data display as detailed in panel E). **H.,** spleen cells from naïve mice and LCMV-immune mice were cultured for 24h under conditions of “withdrawal apoptosis” in the presence of titrated amounts of the FASN inhibitor C75 or vehicle. To account for the differential survival capacity of the different CD8^+^T cell subsets in the absence of inhibitor (O CD8^+^T_M_ > Y CD8^+^T_M_ > CD8^+^T_N_ > CD8^+^T_E_; ***Fig.2G*** and not shown), their relative survival in vehicle cultures was normalized to 100%. Bottom panel: relative survival of indicated CD8^+^T_M_ populations at 30μM C75. **I.,** impact of the LAL inhibitor chloroquine on CD8^+^T cell survival; experimental design as in panel H. Statistical analyses were performed using one-way ANOVA with Dunnett’s multiple comparisons test (panels A, D, E, G and H/I bar diagrams) or Student’s t-test (panel B) comparing indicated CD8^+^T cell populations (the parenthetical asterisk in the upper bar diagram in panel G indicates significance by Student’s t-test but not ANOVA); n≥3 individual mice per group for all experiments conducted independently 2-3 times.

With regard to glucose metabolism, our transcriptional p14 T_M_ data indicated that within the family of facilitative glucose transporters, robust gene expression was restricted to stable *Slc2a1*/Glut1 and progressively declining *Slc2a3*/Glut3 mRNA species (***Fig.4D***). Yet while corresponding Glut3 protein expression levels mirrored the decline of *Slc2a3* mRNA, total Glut1 expression was subject to distinct translational modulations: high in CD8^+^T_E_, reduced in young CD8^+^T_M_ but intermediate in aged CD8^+^T_M_ (***Fig.4E***). Importantly, a specific interrogation of surface Glut1 confirmed the enhanced expression by old *vs.* young CD8^+^T_M_, and the differential Glut1 levels in CD8^+^T_E/M_ populations correlated precisely with their respective glucose uptake capacities (***Fig.4E***). In contrast, greater rates of glucose uptake by CD8^+^T_N_ than either CD8^+^T_E_ or young T_M_, also observed in other reports [44, 45], did not correspond to enhanced Glut1 levels in our experiments (***Fig.4E***); however, neither glucose uptake nor *in vitro* survival of resting T cells is affected by Glut1-deficiency and may instead rely on related transporters such as Glut3 [46]. The notion of enhanced glucose utilization by aged as compared to young CD8^+^T_M_ is further supported by the pattern of CD8^+^T cell-expressed insulin receptor (*Insr*/CD220) that significantly increases with CD8^+^T_M_ age (***Figs.S3A & 4F***). Insulin not only regulates glucose uptake but also acts as a major growth factor that increases protein translation [47]. In fact, of the 26 gene sets demonstrating a progressive enrichment in aging p14 T_M_, nearly half are captured under the general category of “genetic information processing” that includes pathways for transcription; translation; folding, sorting and degradation; as well as replication and repair (***Fig.S3B***).

If CD8^+^T_M_ aging fosters a trend towards increased glucose utilization, it should simultaneously decrease OxPhos and FA utilization, and our GSEAs indicate that this is the case (***Fig.4C***). To directly determine the amount of stored fat in CD8^+^T cells, we quantified neutral lipid content in CD8^+^T_N_ and virus-specific CD8^+^T_E/M_ populations. As expected [45] and albeit subtle, young CD8^+^T_M_ contained fewer neutral lipids than CD8^+^T_E_ but old CD8^+^T_M_ stored even less (***Fig.4G***). In further agreement with O’Sullivan *et al.* [45] we also noted a decreased capacity for long-chain FA (FL C_16_) and low-density lipoprotein (LDL) uptake in young CD8^+^T_M_ as compared to CD8^+^T_E_, a competence that, importantly, eroded even further with age (***Fig.4G***). Reduced FA uptake, however, is not *per se* an indicator for decreased FA metabolism since CD8^+^T_M_ fuel their bioenergetics needs in a “futile cycle” that utilizes extracellular glucose to support both increased FA synthesis (FAS) and FAO [45]. With the aim to delineate the relative contribution of FAS and FAO to CD8^+^T_M_ metabolism [48] specifically in the context of aging, we incubated the various CD8^+^T cell populations in the presence of titrated amounts of selected pharmacological inhibitors and assessed their respective survival. Overall, both young and old CD8^+^T_M_ proved more resistant to inhibition of lipogenesis or lipolysis than either CD8^+^T_E_ or T_N_ (***Figs.4H/I & S3C-E***). Yet subtle differences between young and aged CD8^+^T_M_ could be discerned at particular inhibitor concentrations. Here, inhibition of fatty acid synthase (FASN) by 30μM of the compound C75 compromised survival of young *vs.* old CD8^+^T_M_ to a greater extent suggesting that aged CD8^+^T_M_ are somewhat less reliant on FAS (***Figs.4H & S3C***). Considering the lipolytic machinery of CD8^+^T cells, the recent work by O’Sullivan *et al.* ruled out a role for adipose triglyceride lipase (ATGL) in CD8^+^T_M_ formation and survival [45]; likewise, we found that both young and aged CD8^+^T_M_, in contrast to CD8^+^T_N_ and T_E_, were completely resistant to ATGL inhibition (***Fig.S3D***). Rather, hydrolysis of neutral lipids appears to preferentially rely on lysosomal acid lipase (LAL) [45] and in our experiments, blockade of LAL activity by inhibition of lysosomal acidification with 200μM chloroquine demonstrated a comparatively enhanced death of old CD8^+^T_M_ indicating a greater need for these cells to mobilize FA for FAO (***Figs.4I & S3E***).

Lastly, we wanted to determine how the subtle metabolic alterations in aging CD8^+^T_M_ populations relate to their overall “metabolic fitness”. Here, our determination of mitochondrial mass and membrane potential failed to document consistent differences but in aggregate, we observed a trend towards enhanced fitness by old CD8^+^T_M_ (***Fig.S3F*** and not shown). In support of this assessment, we also note that PGC-1α, a master regulator of mitochondrial biogenesis most recently shown to improve the bioenergetics of LCMV-specific CD8^+^T cells in a chronic infection model [44], is comparatively elevated at both mRNA and protein levels in aged CD8^+^T_M_ (***Fig.S3G/H***). In summary, we conclude that the “mixed metabolic phenotype” of long-term CD8^+^T_M_ populations emerges through a partial reversal of metabolic adaptations that control and accompany the original transition from CD8^+^T_E_ to young CD8^+^T_M_ stage, and that the “intermediate” metabolic profile of old CD8^+^T_M_ likely contributes to their greater recall capacity [1, 5]. Defining a precise inflection point for this “metabolic switch” during CD8^+^T_M_ aging will be difficult given the delicate and only partial nature of metabolic adaptations, but it is well possible that a net effect of these processes may become discernible only at later stages of the extended CD8^+^T_M_ evolution [5].

### Increasing abundance and precipitous maturation of aging CD8^+^T_M_ in the splenic white pulp

The extended maturation of circulating aging CD8^+^T_M_ populations [1] proceeds in the face of their continued anatomic redistribution but without apparent alteration of total CD8^+^T_M_ maintained in various lymphoid organs and NLTs [14, 49-51]. Although there are some exceptions to this rule, *e.g.* the natural decline of influenza virus-specific CD8^+^T_M_ in lung airways and associated loss of immune protection [52], it has remained unclear how exactly the phenotypic conversion of aging CD8^+^T_M_ may modulate their trafficking patterns [53]. The gradual re-expression of CD62L in particular [1, 17] would be expected to affect the anatomical distribution of older CD8^+^T_M_. For example, young p14 T_EM_ and T_CM_ subsets, distinguished according to CD62L expression and with differential sensitivity to the chemokines CCL19 and CXCL12, preferentially localize to splenic red pulp (RP) and white pulp (WP), respectively [54]. The progressive upregulation of CD62L, CCR7 (CCL19 receptor) and CXCR4 (CXCL12 receptor) by aging splenic CD8^+^T_M_ [1], confirmed and extended here to blood-borne CD8^+^T_M_ with different LCMV specificities (***Fig.5A*** and not shown), may therefore also promote an altered positioning of these cells within the spleen. To evaluate this possibility, we employed the i.v. injection of fluorochrome-conjugated CD8β antibody that readily labels CD8^+^T cells found in vascular contiguous compartments (including RP) but not tissue stroma and parenchyma (including WP) [55, 56] (***Fig.S4A***). While the total number of specific CD8^+^T_M_ in the spleen does not change over time [14], their differentiation according to RP/WP residence demonstrated a pronounced increase from ∼15% to ∼60% in the WP of aging mice (***Fig.5B***). A concurrent phenotypic stratification of RP/WP subsets according to markers that are substantially up- or down-regulated by aging CD8^+^T_M_ [1] further revealed striking differences in young mice: the ∼15% of young D^b^NP_396_^+^ CD8^+^T_M_ residing in the WP, despite preserving some phenotypic heterogeneity, for the most part already adopted properties comparable to aged CD8^+^T_M_ (CD27^hi^, CD62L^hi^, CD127^hi^, CXCR3^+^, CD43^lo^, KLRG1^-^, CX3CR1^lo^) whereas RP cells (representing ∼85% of splenic D^b^NP_396_^+^ CD8^+^T_M_) exhibited a contrasting and largely “immature” phenotype (***Fig.5C-E***); these differences also pertained to more subtle aspects of CD8^+^T_M_ aging such as SSC properties and CD8α expression levels (albeit not cellular size) (***Fig.5E***). In aged LCMV-immune mice, and in agreement with the observation that phenotypic maturation affects both splenic and blood-borne CD8^+^T_M_ (ref.[1] and ***Figs.3A/B & 5A***), the dissimilarity of WP and RP D^b^NP_396_^+^ CD8^+^T_M_ mostly disappeared and both populations presented with an aged phenotype (though the RP subset retained somewhat elevated CD43, KLRG1 and CX3CR1 expression) (***Fig.5C-E***). Nearly identical results were also obtained for young and old D^b^GP_33_^+^ CD8^+^T_M_ in splenic RP/WP compartments (***Fig.S4B-D***). Lastly, a direct comparison of young and old CD8^+^T_M_ in the RP confirmed their marked phenotypic differences but the WP subsets, to a lesser degree, also demonstrated evidence for further age-associated phenotype maturation (***Fig.S4E***). Altogether, these observations reveal the gradual emergence of co-regulated complex CD8^+^T_M_ phenotypes as well as their distinct spatiotemporal segregation that accompanies the more global architectural changes recently reported for the aging murine spleen [57].

**Figure 5.**
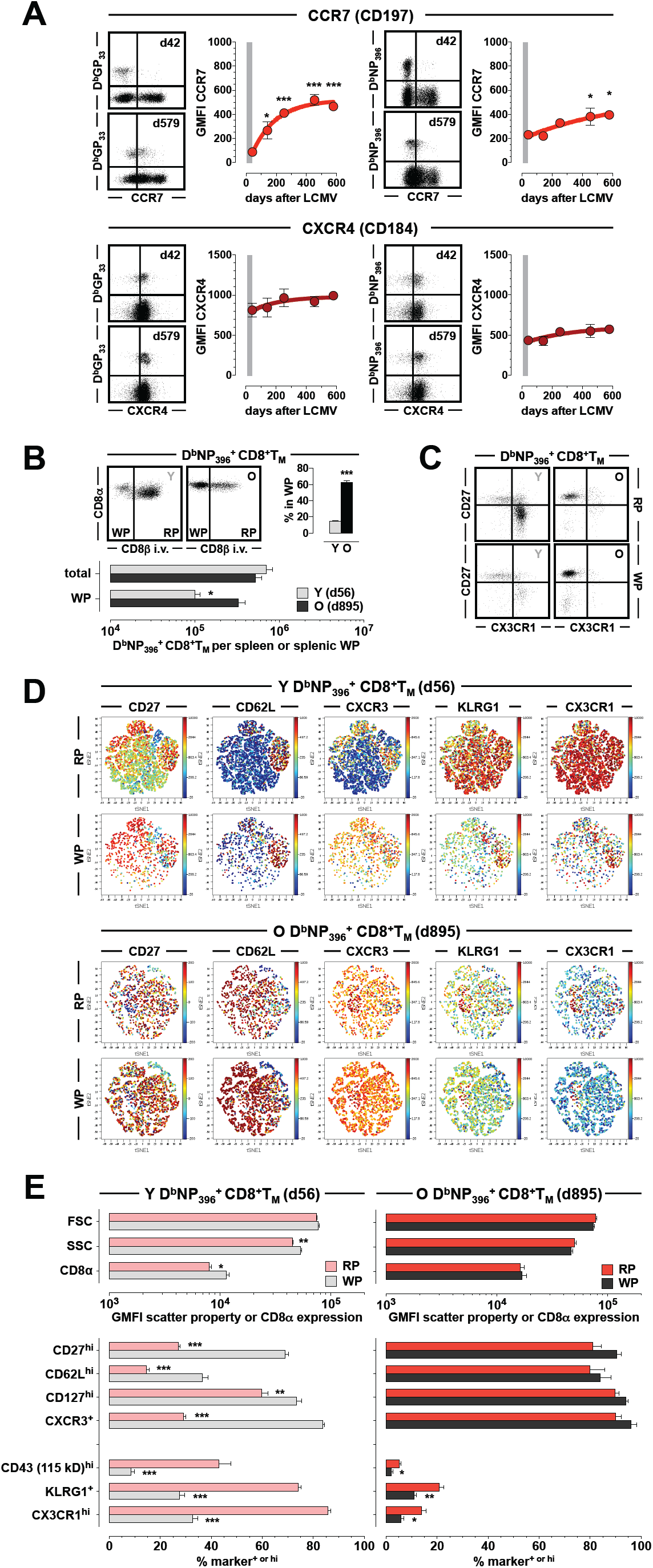
Increasing abundance and accelerated maturation of aging CD8^+^T_M_ in the splenic WP. **A.,** temporal regulation of CCR7 (top) and CXCR4 (bottom) expression by D^b^GP_33_^+^ (left) and D^b^NP_396_^+^ (right) CD8^+^T_M_ in peripheral blood; dot plots are gated on CD8^+^T cells and CXCR4 expression was revealed by intracellular stains (n=4 mice/time point). Although the subtle increase of CD8^+^T_M_-expressed CXCR4 is not statistically significant in the present data sets, the trend is apparent and in agreement with significant differences shown in related experiments (***Fig.S5A*** and ref.[1]). **B.,** relative abundance of D^b^NP_396_^+^ CD8^+^T_M_ in the splenic WP of young and aged mice as revealed by intravascular CD8 staining. **C.,** phenotypic properties of young and old D^b^NP_396_^+^ CD8^+^T_M_ in splenic RP *vs.* WP. **D.,** viSNE rendering of the D^b^NP_396_^+^ CD8^+^T_M_ phenotype space in RP *vs.* WP of young (top) and old (bottom) LCMV-immune mice. **E.,** individual phenotypic characteristics of D^b^NP_396_^+^ CD8^+^T_M_ RP and WP populations in young (left) and old (right) mice (panels B-E: n≥3 mice/time point analyzed in 2 separate experiments; for further details on intravascular staining and viSNE analyses, see Methods).

### Progressive accumulation of aging CD8^+^T_M_ in peripheral lymph nodes

Another potential consequence of increasing CD62L, CCR7 and/or CXCR4 expression by aging CD8^+^T_M_ (***Fig.5A***) is the gradual acquisition of an enhanced LN tropism [58, 59], especially since earlier trafficking studies have demonstrated the unequivocal requirement for virus-specific CD8^+^T_M_-expressed CD62L [60] and chemokine receptors [50] to enter LNs under steady-state conditions. Indeed, a first suggestion in support of this conjecture has come from a recent study that reported a greater proportion of “late” p14 T_M_ as compared to “early” p14 T_M_ in inguinal LNs [5]. To examine if the “LN-homing phenotype” of aged CD8^+^T_M_ confers a preferential redistribution to peripheral LNs *at large*, we enumerated specific CD8^+^T_M_ in young and old LCMV-immune mice. In the absence of age-associated changes in LN cellularity, we observed an up to 10-fold increase of specific CD8^+^T_M_ frequencies and numbers in aged mice (***Fig.6A-C***), and a longitudinal analysis of mesenteric LNs (MLN) revealed a slow and continuous accumulation of CD8^+^T_M_ with an estimated population doubling time of ∼190 days (***Fig.6D***). We next assessed the capacity of aging CD8^+^T_M_ to enter peripheral LNs by performing a competitive homing experiment (***Fig.6E***). In brief, p14 T_M_ were enriched from young and old LCMV-immune p14 chimeras, differentially labeled with CFSE, combined at a ratio of 1:1, and transferred into naïve B6 recipients. Upon retrieval 48h later, this ratio was skewed to >10:1 in favor of old p14 T_M_ in peripheral LNs but not blood or spleen demonstrating that aging CD8^+^T_M_ in fact acquire a capacity for facilitated LN access (***Fig.6E***).

**Figure 6.**
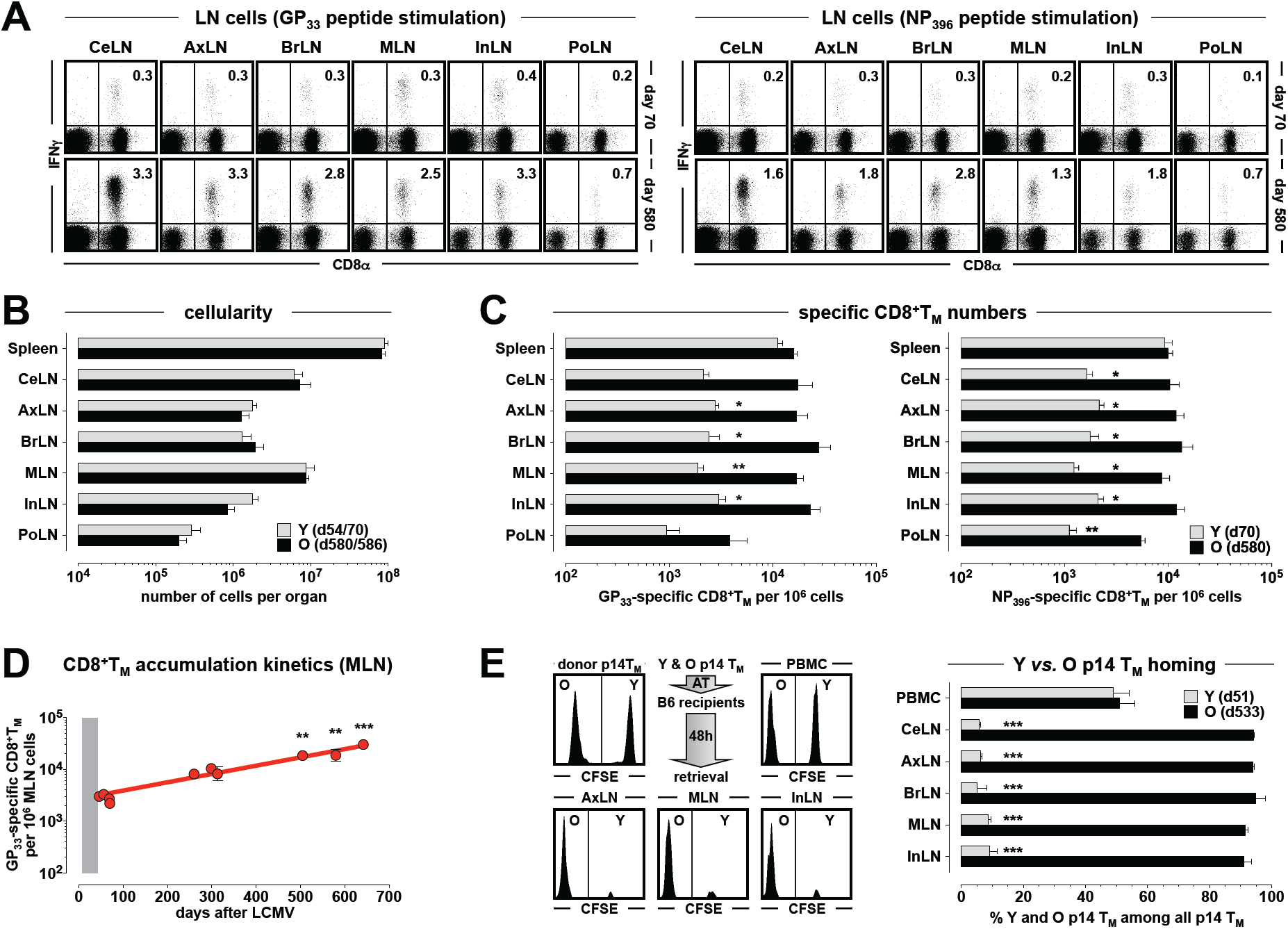
Progressive accumulation of aging CD8^+^T_M_ in peripheral LNs. **A.,** LNs were harvested from young and old LCMV-immune B6 mice, restimulated with GP_33_ (left) or NP_396_ (right) peptides and stained for CD8α and intracellular IFNγ. Values indicate frequencies of epitope-specific CD8^+^T_M_ among all LN cells (similar results were obtained for CD8^+^T_M_ specific for the subdominant GP_276_ epitope, not shown). CeLN: cervical LN, AxLN: axillary LN, BrLN: brachial LN, MLN: mesenteric LN, InLN: Inguinal LN, PoLN: popliteal LN. **B.,** cellularity of spleen and indicated LNs obtained from young and old LCMV-immune B6 mice. **C.,** numbers of GP_33_-(left) and NP_396_-specific (right) CD8^+^T_M_ in spleen and peripheral LNs of young and old mice (n=3; data from 1/4 independent experiments). **D.,** progressive accumulation of GP_33_-specific CD8^+^T_M_ in the MLN of aging LCMV-immune mice (n=2-4 for each time point, asterisks indicate statistical significance comparing young [∼d50] and older mice). Comparative non-linear regression analyses for the period from ∼d50-d650 revealed a best curve fit using an exponential growth model (r^2^=0.88) and thus permitted the calculation of a population doubling time of t_D_=188 days. **E.,** splenic p14 T_M_ populations enriched from young (d51) and old (d533) p14 chimeras were differentially labeled with CFSE, combined at a ratio of 1:1 (upper left histogram), transferred i.v. into B6 recipients and retrieved 48 hours later from various tissues (experimental flow chart and other histograms); the bar diagram summarizes the relative composition of young and old p14 T_M_ populations recovered from blood and indicated LNs (representative data from 1/2 similar experiments).

### Increased CD62L expression promotes improved LN access for aging CD8^+^T_M_

Similar to polyclonal CD8^+^T_M_ (***Fig.5A*** and ref.[1]), old p14 T_M_ exhibited higher expression levels of CCR7, CXCR4 and in particular CD62L (***Figs.S5A & 7A***). To determine if CD62L contributed directly to the facilitated LN access of aged CD8^+^T_M_, we conducted an *in vivo* homing assay with old p14 T_M_ under conditions of CD62L blockade and observed an 82-93% reduction of p14 T_M_ accumulation in peripheral LNs (***Fig.7A***). Similar experiments designed to evaluate the role of chemokine receptors by pretreatment of young and old donor p14 T_M_ with pertussis toxin (Ptx) revealed, as expected [50], a profound inhibition of p14 T_M_ trafficking to LNs (***Fig.S5B***). The relative reduction, however, appeared especially pronounced for young p14 T_M_ indicating a slight advantage for aged p14 T_M_ to utilize Ptx-insensitive pathways for residual LN access (***Fig.S5B***). The importance of CD62L in conveying an enhanced LN tropism to CD8^+^T_M_ populations was further illustrated by use of the “virus titration chimeras” discussed above. Following infection of p14 chimeras with escalating titers of LCMV and generation of T cell memory 7 weeks later, p14 T_M_ expression of CD62L but not CCR7 or CXCR4 significantly declined as a function of increasing viral challenge dosage (***Fig.7B*** and data not shown), and reduced CD62L expression correlated with an impaired accumulation of p14 T_M_ in peripheral LNs (***Fig.7B***). Thus, the LN tropism of CD8^+^T_M_, in addition to their survival/Bcl-2 expression (***Fig.2J***), multiple phenotypic and functional properties, and their II^o^ reactivity [1], can be experimentally controlled in a fashion that accelerates or delays the CD8^+^T_M_ maturation process *at large*.

**Figure 7.**
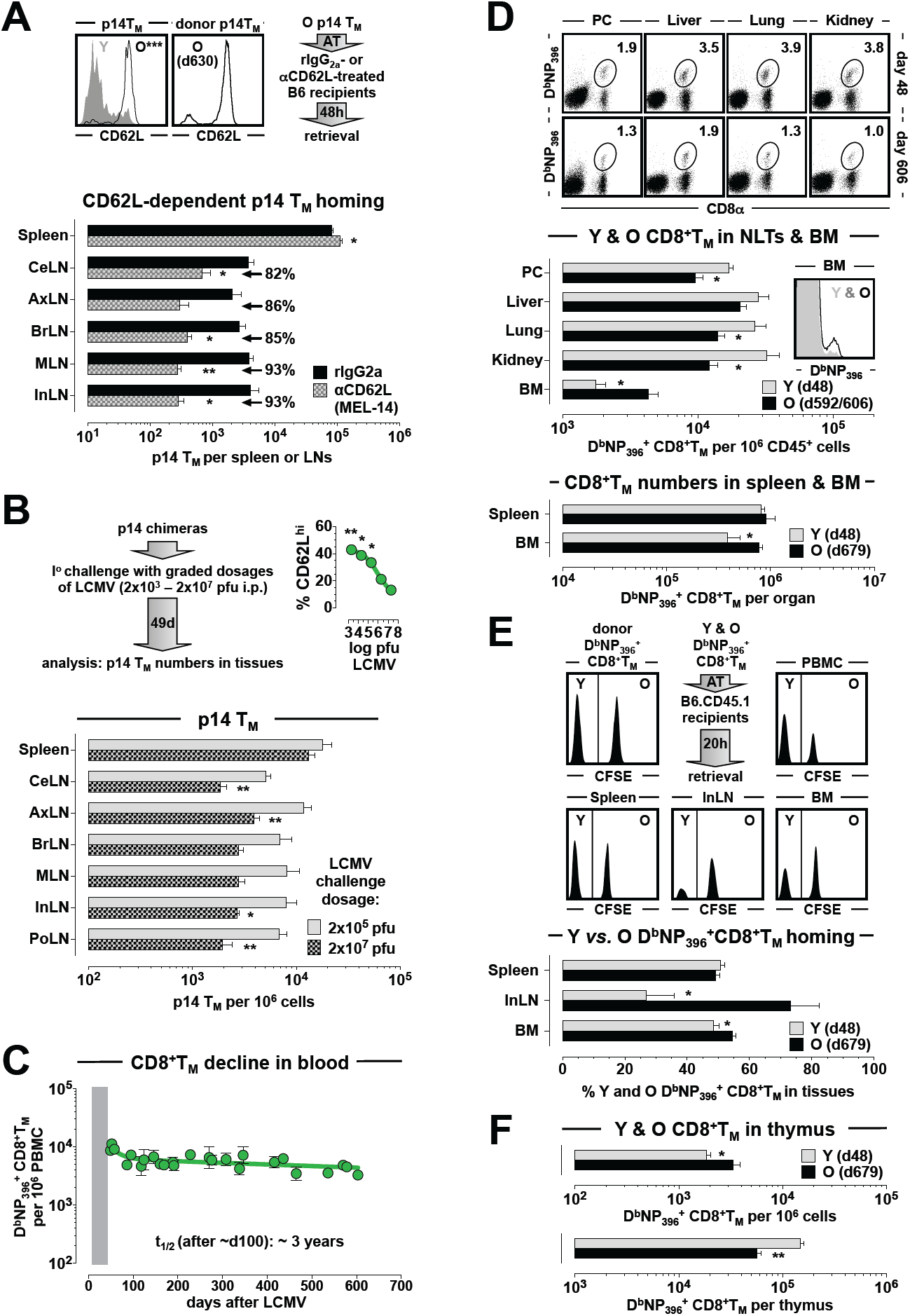
Redistribution of aging CD8^+^T_M_ from NLTs to lymphoid tissues. **A.,** upper left/middle: CD62L expression of young and aged p14 T_M_ used for homing assays in ***Fig.6E*** (asterisks indicate significant differences with n=4-5 mice), and of old donor p14 T_M_ used for CD62L blocking studies. Upper right: experimental flow chart for p14 T_M_ trafficking experiments. Bottom: enumeration of p14 T_M_ in spleen and LNs of recipient mice treated with αCD62L or control antibodies; the values indicate the extent of reduced LN trafficking as a consequence of CD62L blockade. **B.,** left: experimental flow chart depicting the generation of “virus titration chimeras”; right: CD62L expression levels of p14 T_M_ (d49) as a function of original virus challenge dosage. Bottom enumeration of p14 T_M_ in spleen and LNs of LCMV-immune p14 chimeras infected with 2×10^5^ or 2×10^7^ pfu LCMV. **C.,** subtle decline of aging D^b^NP_396_^+^ CD8^+^T_M_ in peripheral blood (combined data from multiple independent experiments); the theoretical population half-life beyond d100 after infection was calculated to be ∼3 years. **D.,** quantification of D^b^NP_396_^+^ CD8^+^T_M_ isolated from lymphatic and nonlymphoid tissues of young and old LCMV-immune B6 mice. Dot plots and histograms are normalized to display 1.7×10^4^ CD45^+^ cells with values indicating the fraction of D^b^NP_396_^+^ CD8^+^T_M_; the bar diagrams display representative results from two independent experiments. **E.,** homing of young and old D^b^NP_396_^+^ CD8^+^T_M_ was assessed by differential CFSE labeling of donor populations, combination at a ratio of 1:1 (upper left histogram), i.v. transfer of 5.5×10^4^ D^b^NP_396_^+^ CD8^+^T_M_ each into B6.CD45.1 recipients, and retrieval from indicated tissues 20 hours later. **F.,** enumeration of young and old D^b^NP_396_^+^ CD8^+^T_M_ in the thymus; n≥3 individual mice per group for all experiments.

### Loss of aging CD8^+^T_M_ from peripheral blood and nonlymphoid tissues

Based on the above evidence, and in the absence of locally increased homeostatic proliferation (***Fig.3E/F***), the progressive accumulation of aging CD8^+^T_M_ in secondary lymphoid tissues (***Figs.5B, S4B & 6***) most likely emerged through the redistribution of CD8^+^T_M_ from other anatomic reservoirs. We estimated, according to the numbers of specific CD8^+^T_M_ in the LNs of young and aged LCMV-immune mice (***Fig.6A-C***) as well as the number and variable size of murine LNs [61], that over a period of ∼17 months, up to 1×10^6^ NP_396_ - and 1.9×10^6^ GP_33_-specific CD8^+^T_M_ were added to the entire LN pool. Given the stable CD8^+^T_M_ numbers in the spleen [14], the potential sources for the new LN CD8^+^T_M_ are therefore the blood and marginated pool (BMP), as well as NLTs. In a most recent and comprehensive accounting of organism-wide CD8^+^T_M_ distribution, based on an evaluation of LCMV-immune p14 chimeras, Steinert *et al.* demonstrated that NLTs and BMP (excluding splenic red pulp) together contain ∼6×10^6^ p14 T_M_ [56]. The p14 model used therein and our B6 system are roughly comparable since flow cytometry-based calculations revealed the presence of ∼2.9×10^6^ splenic p14 T_M_ while we documented a total of ∼2.0×10^6^ endogenously generated NP_396_/GP_33_-specific CD8^+^T_M_ in the spleen (***Fig.6B/C*** and not shown). In regards to LN-residing CD8^+^T_M_, however, the models are expected to differ due to increased p14 T_N_ numbers used for chimera construction [56], correspondingly accelerated upregulation of CD62L by p14 T_M_ [1], and an experimental evaluation at a somewhat later time points (4-5 months after challenge) [56] that together should result in apparently enhanced LN accumulation. Indeed, the reported grand total of ∼2.3×10^6^ p14T_M_ in peripheral LNs [56] clearly exceeded the ∼4.3×10^5^ NP_396_/GP_33_-specific CD8^+^T_M_ we found in the LN compartment of B6 mice at ∼2 months following LCMV infection (a 5.4-fold difference). With these caveats in mind, we calculated that in the time of ∼2-19 months after infection, a less than 2-fold loss from BMP and NLTs could account for the corresponding gain of NP_396_/GP_33_-specific CD8^+^T_M_ in the LNs of old LCMV-immune B6 mice.

To test this prediction, we first evaluated the preservation of D^b^NP_396_ CD8^+^T_M_ in the blood by combining data obtained in numerous experiments performed over a period of several years. Interestingly, the aggregate data uncovered an unexpected biphasic loss of blood-borne CD8^+^T_M_ (***Fig.7C***). In the period of ∼7-14 weeks after virus challenge, and thus well after completion of the “contraction phase” in the spleen [14], specific CD8^+^T_M_ numbers continued to decline in the blood before attaining seemingly stable levels around day 100 after infection. A careful inspection of subsequent time points, however, revealed a subtle decrease of blood-borne CD8^+^T_M_ with a theoretical population half life of ∼3 years (***Fig.7C***). This finding is noteworthy since it evokes, even under experimental conditions that optimize CD8^+^T_M_ preservation, the natural decline of blood-borne virus-specific CD8^+^T_M_ in humans [12]. In as much as the cumulative ∼60% loss (between weeks 7 and 86) of specific CD8^+^T_M_ from peripheral blood also reflects a changing CD8^+^T_M_ abundance in the larger BMP, these cells could provide a relevant contribution to the growing CD8^+^T_M_ LN pool. The biphasic erosion of blood-borne CD8^+^T_M_ (***Fig.7C***), however, would seem at odds with the dynamics of CD8^+^T_M_ accumulation in the LNs (***Fig.6D***). We therefore proceeded with an enumeration of young and old CD8^+^T_M_ in NLTs (peritoneal cavity, liver, lung, kidney) and observed a 1.4-to 2.7-fold relative reduction of aged CD8^+^T_M_ numbers (***Fig.7D***). Thus, both theoretical considerations and experimental results support the notion that a loss of aging CD8^+^T_M_ is not restricted to the lung [5, 52] but involves the BMP and especially NLTs in general.

### CD8^+^T_M_ trafficking and the “tissue resident memory T cell (T_RM_)” paradigm

How can the above conclusions be reconciled with the notion that NLTs are preferentially populated by non-recirculating T_RM_ [62]? According to Steinert *et al.*, ∼9% of CD8^+^T_M_ found in NLTs can in fact recirculate, a fraction that is lower in some (*e.g.,* lung) but higher in other (*e.g.,* liver) compartments [56]. These calculations are based on parabiosis experiments that were conducted, similar to multiple other studies, over a period of just ∼1 month [56]. A notably longer observation period was employed by Jiang *et al.* who found that the frequencies of skin CD8^+^T_RM_ in the donor parabiont declined by ∼2-fold between 8 and 24 weeks after surgery suggesting limits to CD8^+^T_RM_ longevity and/or mobilization of the CD8^+^T_RM_ compartment [63]. The latter observation is not only in agreement with a classic study that reported a trend towards continued equilibration of CD8^+^T_RM_ within intestinal lamina propria and epithelium for at least 8 weeks [50] but also consistent with our experiments that compare CD8^+^T_M_ populations recovered from NLTs at time points separated by ∼18 months and thus may offer sufficient time for some CD8^+^T_RM_ to re-enter the circulation. Of further importance is the recent observation that traditional flow cytometry-based methods of CD8^+^T_M_ quantification in NLTs markedly underestimate the true number of CD8^+^T_M_ found in these tissues [56]. While our quantification of young and old CD8^+^T_M_ in liver, lung and kidney therefore cannot accurately account for absolute CD8^+^T_M_ numbers, it is the *relative* reduction of CD8^+^T_M_ recovered from the NLTs of aged LCMV-immune animals, readily revealed even by use of flow cytometry, that is important for the present context. Consistent with this interpretation, we also observed an age-associated decrease of CD8^+^T_M_ numbers in the peritoneal cavity (***Fig.7D***), an organ that is not subject to the inefficiency of CD8^+^T_M_ recovery from solid NLTs. Finally, in considering the role of CD8^+^T_RM_ as highly effective first responders to infections re-encountered at body surfaces [64] and the established role of LN-residing CD8^+^T_M_ as direct precursors for II^o^ CD8^+^T_E_ expansions, it is worth noting that LN CD8^+^T_M_ themselves also act as “gate-keepers” and immediate effectors capable of curtailing peripheral infections and preventing systemic viral spread [65]. In fact, following a footpad LCMV challenge of mice that received limiting numbers of young *vs.* old CD8^+^T_M_, we found that only the latter population prevented systemic dissemination of the virus (not shown). Therefore, the gradual accumulation of aging CD8^+^T_M_ in peripheral LNs, even at the expense of CD8^+^T_M_ in NLTs, may represent a progressively enhanced “strategic positioning” in anatomic locations that constitute a critical site for both local pathogen control and the coordination of effective recall expansions [66].

### Increased accumulation of old CD8^+^T_M_ in primary lymphatic tissues

Considering the tissue redistribution of aged CD8^+^T_M_ in their overall numerical context (***Figs.5B, 6 & 7C/D***), it appears that the relative loss from NLTs and blood might even exceed the corresponding gain in peripheral LNs. A clue to another anatomic site for potential CD8^+^T_M_ accrual comes from the increased CXCR4 expression by old antiviral CD8^+^T_M_ (***Figs.5A, S5A*** & ref.[1]). CXCR4 is held to be a “BM homing receptor” and consistent with this notion, recent work demonstrated that conditional CXCR4 deletion in LCMV-specific T cells resulted in a reduced abundance of CD8^+^T_M_ populations especially in the BM [67]. Thus, it is conceivable that greater CXCR4 expression levels by aged CD8^+^T_M_ preferentially promote increased BM access, an important anatomic niche for CD8^+^T_M_ [68]. Indeed, the frequencies and numbers of D^b^NP_396_^+^ and D^b^GP_33_^+^ CD8^+^T_M_ retrieved from the BM of aging LCMV-immune mice roughly doubled over a period of ∼1.5 years (***Fig.7D***) though in contrast to LNs, accumulation of aging CD8^+^T_M_ in the BM was independent of CD62L (***Fig.S5C***). In competitive homing experiments similar to those shown in ***Fig.6E*** but conducted here with endogenously generated CD8^+^T_M_, aged CD8^+^T_M_ also displayed a slightly enhanced BM tropism; at the same time, their facilitated LN access was expectedly more pronounced (***Fig.7E***).

Finally, the apparently generalized pattern of age-associated increasing CD8^+^T_M_ abundance in both secondary (splenic WP, LN) and primary (BM) lymphatic tissues also warranted an analysis of the thymus. Interestingly, we observed an almost 2-fold *relative* increase of old over young CD8^+^T_M_ populations for this primary lymphatic organ (counts normalized to 10^6^ cells); due to thymic involution, however, absolute numbers of aged CD8^+^T_M_ were expectedly reduced, here by a factor of ∼2.5 (***Fig.7F***).

## CONCLUSIONS

As detailed in our recent work on CD8^+^T cell memory [1], aging of established antiviral CD8^+^T_M_ populations introduces a series of cumulative molecular, phenotypic and functional changes that collectively confer naïve-like T cell traits, greater proliferative potential and protective capacities onto old CD8^+^T_M_ populations. To account for these sweeping processes in a simple fashion, we have introduced the “rebound model” of CD8^+^T_M_ maturation according to which the extent of initial CD8^+^T_E_ differentiation directly determines the kinetics of protracted CD8^+^T_M_ “de-differentiation” [1]. We now demonstrate that this remodeling process also impinges on the homeostasis of CD8^+^T_M_ as evidenced by their evolving survival capacity, metabolic adaptations and microanatomic redistribution. Here, both the Bcl-2-dependent enhancement of apoptosis resistance and the accumulation of old CD8^+^T_M_ in lymphoid tissues (including the CD62L-guided peripheral LN access/residence) as a likely consequence of a redistribution from NLTs and blood are consistent with the progressive modulation of aging CD8^+^T_M_ phenotypes, in particular at the level of increasing CD62L, CD122, CD127, CCR7 and CXCR4 expression [1-3, 5, 17]. We further document that the gradual acquisition of mature phenotypes by aging CD8^+^T_M_ populations proceeds through co-regulated modulation of receptor/ligand expression and at a pace that is contingent on the specific microenvironment (i.e., accelerated in splenic WP, delayed in RP). In addition, all of these dynamics are readily captured by the basic tenet of the “rebound model” that posits a broad harmonization of CD8^+^T_M_ and T_N_ traits while simultaneously reinforcing the development of a simple CD8^+^T_M_ core signature [1].

The imperviousness of aging CD8^+^T_M_ to changes of their basal homeostatic proliferation rates, however, was unexpected. Our results document a simple association between cytokine receptor (CD122/CD127) expression levels and functionality, and the importance of CD127 abundance as well as the intermittent rather than continuous IL-7 signaling for the homeostasis of naïve CD8^+^T cell populations has been illustrated by the work of A. Singer’s group [69]. Yet we previously noted a lack of association between CD127/CD122 expression levels on CD8^+^T_M_ and their tissue-specific pace of homeostatic turnover [38], and the heightened responsiveness of aged CD8^+^T_M_ to IL-7 and IL-15 as shown here failed to confer increased homeostatic proliferation rates. Old CD8^+^T_M_ may therefore have adopted an exquisite balance with age-associated changes in various tissue microenvironments; the homeostasis of CD8^+^T_N_ and pathogen-specific CD8^+^T_M_, though reliant on the same cytokines (IL-7, IL-15), may be regulated in a differential manner; or other factors contributing to the regulation of CD8^+^T_M_ homeostasis may become more dominant over time. We also note that changing levels of CD8^+^T_M_-expressed CXCR4, recently proposed to control the homeostatic turnover of CD8^+^T_M_ [67], had no apparent impact on their homeostatic self-renewal over time. A recent analysis of human CD45RO^+^CD8^+^T_MP_ populations also found no differences in homeostatic turnover rates between young and healthy elderly individuals [70].

Though subtle, the metabolic adaptations of aging CD8^+^T_M_ would appear to contradict the “rebound model” since they are characterized by a partial re-acquisition of CD8^+^T_E_-like profiles, in particular an increase of glucose utilization [40]. Yet the shift towards enhanced glucose uptake, decreased neutral lipid content as well as reduced FA and LDL uptake also indicates a gradual return, albeit incomplete, towards respective CD8^+^T_N_ capacities. Nevertheless, CD8^+^T_N_ consistently displayed greater sensitivity to *in vitro* FAS and FAO inhibition than either young or old CD8^+^T_M_ suggesting that the latter cells’ distinctive and evolving metabolic profiles should be considered part of the memory “core signature” that distinguishes CD8^+^T_M_ from T_N_.

Three aspects of CD8^+^T_M_ homeostasis will require further clarification to define relevant age-associated adaptations and their potential impact on II^o^ reactivity and immune protection in more detail. 1., the progressive conversion of aging CD8^+^T_M_ documented primarily for spleen and blood [1-3, 5] will have to be considered for other tissues [62], in the context of continued CD8^+^T_M_ subset migration *vs.* extended tissue residence (including the precise developmental relations and potential phenotypic/functional modulation of CD8^+^T_M_ populations as they enter and exit various tissues) [38, 71, 72], and for human CD8^+^T_M_ [73]. 2., the transcriptional control of CD8^+^T_M_ aging is another topic of broad interest. For example, among the major transcriptional regulators of CD8^+^T_E/M_ differentiation predicted on the basis co-regulated gene expression in activated CD8^+^T cells [74] are several TFs (*Tcf4*, *Zeb2*, *Rora*, *Hif1a*, *Arntl*) that also demonstrate progressive downmodulation in aging CD8^+^T_M_ [1]. In agreement with this observation, enhanced activity of hypoxia-inducible factors (HIFs) was recently shown to sustain a CD8^+^T_E_-like state [75] while *Zeb2*-deficiency accelerated CD8^+^T_CM_ formation [76]; the extent to which the evolution of complex TF expression profiles in aging CD8^+^T_M_ supports a return to a CD8^+^T_N_-like transcriptional program while simultaneously reinforcing the emergence of a highly focused CD8^+^T_M_ “core signature” is currently under investigation. 3., in conjunction with transcriptional regulation, epigenetic DNA and chromatin modifications provide irreducible contributions to the specification of CD8^+^T_M_ fates [77]. Though it remains unclear if established CD8^+^T_M_ are subject to epigenetic modulations under steady-state conditions, it is conceivable that exposure to or withdrawal from different microenvironmental cues may alter the epigenetic landscape of aging CD8^+^T_M_.

In summary, the present work confirms and expands the central tenets of the “rebound model” [1] by documenting the fundamentally temporal nature of CD8^+^T_M_ homeostasis and identifying associated determinants for improved CD8^+^T_M_ survival, metabolic alterations and lymphoid tissue homing that collectively brace aged CD8^+^T_M_ for enhanced II^o^ expansion and immune protection. The dynamic adaptations of long-term CD8^+^T cell memory and the possibility to accelerate or delay these processes *at large* [1] provides an experimental framework for the focused interrogation of suitable targets that may be exploited for the prophylactic or therapeutic modulation of specific CD8^+^T_M_ responses. To this end, we have explored elsewhere the specific contribution of 16 molecular pathways to the improved II^o^ reactivity of aged CD8^+^T_M_ populations (manuscripts in preparation).

## METHODS

### Ethics statement

All procedures involving laboratory animals were conducted in accordance with recommendations in the “Guide for the Care and Use of Laboratory Animals of the National Institutes of Health”, the protocols were approved by the Institutional Animal Care and Use Committees (IACUC) of the University of Colorado (permit numbers 70205604[05]1F, 70205607[05]4F and B-70210[05]1E) and Icahn School of Medicine at Mount Sinai (IACUC-2014-0170), and all efforts were made to minimize suffering of animals.

### Mice, virus and challenge protocols

C57BL6/J (B6), congenic B6.CD90.1 (B6.PL-*Thy1*^*a*^/CyJ) and congenic B6.CD45.1 (B6.SJL-*Ptprc*^*a*^ *Pepc*^*b*^/BoyJ) mice were purchased from The Jackson Laboratory; p14 TCRtg mice were obtained on a B6.CD90.1 background from Dr. M. Oldstone (CD8^+^T cells from these mice [“p14 cells”] are specific for the dominant LCMV-GP_33-41_ determinant restricted by D^b^). We only used male mice in this study to avoid potential artifacts that may arise in gender mismatched adoptive transfer settings. LCMV Armstrong (clone 53b) was obtained from Dr. M. Oldstone and stocks prepared by a single passage on BHK-21 cells; plaque assays for determination of virus titers were performed as described/referenced [14]. For I^o^ challenges, 8-10 week old mice were infected with a single intraperitoneal (i.p.) dose of 2×10^5^ pfu LCMV Armstrong; for II^o^ challenges, naïve recipients (aged 8-10 weeks) of various CD8^+^T_M_ populations were inoculated with 2×10^5^ pfu LCMV Arm i.p. All mice were housed under SPF conditions and monitored for up to ∼2 years. Aging LCMV-immune mice were excluded from our study if they presented with 1., gross physical abnormalities such as lesions, emaciation and/or weight loss, 2., lymphatic tumors as indicated by enlarged LNs at time of necropsy or 3., T cell clonal expansions within the virus-specific CD8^+^T_M_ compartment (D^b^NP_396_^+^, D^b^GP_33_^+^ or D^b^GP_276_^+^). According to these criteria, up to ∼30% of aging mice were excluded from the study.

### Tissue processing, cell purification and adoptive transfers (AT)

Lymphocytes were obtained from blood, spleen, lymph nodes (LNs), thymus, peritoneal cavity and bone marrow (BM) according to standard procedures; for an estimate of total BM cells, the content from one femur was multiplied with a coefficient of 15.8 [38] (***Fig.7D***). For isolation of lymphocytes from solid NLTs (liver, lung, kidney), terminally anesthetized mice were sacrificed by total body perfusion with PBS and subsequent organ processing and gradient centrifugation as described [38]. Enrichment of splenic T cells was performed with magnetic beads using variations and adaptations of established protocols. 1., for construction of p14 chimeras [1], p14 T_N_ (CD90.1^+^) were enriched from spleens of naïve p14 mice by negative selection (EasySep Mouse CD8^+^T Cell Enrichment Kit, StemCell Technologies) and transferred i.v. into B6 recipients at indicated numbers prior to LCMV infection 2-24h later (***Fig.2J:*** 2×10^2^–2×10^5^ or 1×10^4^; ***Figs.3C, S2C & S5A/B:*** 5×10^4^; ***Figs.6E, 7A & S5C:*** 2×10^3^, ***Fig.7B:*** 1×10^4^). 2., purification of p14 T_E/M_ for microarray analyses is described in ref.[1]. 3., enrichment of CD8^+^T_M_ from LCMV-immune B6 and B6-congenic donors was performed by depletion of B220^+^ cells (Miltenyi, Invitrogen/Dynal or StemCell Technologies) followed by 1:1 combination at the level of D^b^NP_396_^+^ CD8^+^T_M_, i.v. AT of mixed populations containing 2×10^3^ D^b^NP_396_^+^ congenic CD8^+^T_M_ each into naïve congenic recipients, and challenge with LCMV (***Fig.2H***).

### Flow cytometry

All reagents and materials used for analytical flow cytometry are summarized in ***Table S1***, and our basic staining protocols are described and/or referenced in ref.[1]; in some cases, expression levels were normalized by dividing the GMFI of experimental by the GMFI of isotype control stains (***Fig.2I***). Additional methodologies employed here include the use of various fluorescent dyes/probes (PI, 7AAD, YO-PRO-1, Zombie dyes, DiOC_6_(3), dihydroethidium [HE], Alm Alx488, ThiolTracker Violet, Glut1.RBD.GFP [stained at 37^o^C for detection of surface Glut1]), Mito Tracker Green [MTG], tetramethylrhodamine [TMRE] and JC-1 dye according to manufacturer recommendations and/or published protocols [21, 78, 79] (***Figs.2, 4E, S3F*** and not shown), and the detection of certain intracellular antigens using methanol permeabilization (pSTAT5, Glut1, Glut3) as described [80] or the ebioscience Foxp3/TF buffer set (PGC-1α) (***Figs.3C/D, 4E, S2A & S3H***). Lipid content and lipid/glucose uptake were determined by incubation with Bodipy 493/503 (0.5μg/ml PBS, 10min. at RT) or 37^o^C culture in complete RPMI in the presence of Bodipy FL C16 (overnight at 0.5μg/ml), Bodipy LDL (30min. at 10μg/ml), or 2-NBDG (2h at 100μg/ml) prior to cell surface stains and acquisition (***Fig.4E/G***). Intravascular staining of CD8^+^T cells was adapted from the methodology developed by Anderson *et al.* [55] (i.v. injection of 4μg anti-CD8β-PE [53-5.8] followed by euthanasia 4-5 min later, tissue harvesting/processing and staining with anti-CD8α-BV421 or -PerCP-Cy5.5 [53-6.7], other cell surface receptor/ligand antibodies and MHC-I tetramers; ***Figs.5B-E & S4***). Samples were acquired on FACSCalibur, Accuri C6, Canto, LSRII or LSR Fortessa X-20 flow cytometers (BDBiosciences) and analyzed with DIVA (BDBiosciences) and/or FlowJo (TreeStar) software; dimensionality reduction and data display for polychromatic flow cytometry was performed using the Cytobank platform and the t-SNE algorithm viSNE [81] (input parameters: FSC/SSC properties and CD8α, CD8β, CD27, CD43 (S7), CD62L, CD127, KLRG1, CXCR3, CX3CR1 mean expression levels of young or old D^b^NP_396_^+^ and D^b^GP_33_^+^ CD8^+^T_M_ populations).

### Microarray analyses and qRT-PCR

Details for microarray analyses of highly purified p14 T_E/M_ populations are found in ref.[1], and selected data are shown here in ***Figs.1C/D, 4A/D, S1 & S3A/G***. Gene set enrichment analyses (GSEA) were performed based on filtered data sets obtained for aging p14 T_M_ (d46, d156, d286 and d400) [1] against 186 KEGG gene sets/pathways (http://software.broadinstitute.org/gsea) (***Figs.1A, 2E, 4C, S2B & S3B***). We treated time series as continuous phenotypes and used Pearson’s correlation to determine ranks for each gene. Enrichment scores (ES) were obtained as the maximum deviation from zero of *P*_hit_ – *P*_miss_, where *P*_hit_ and *P*_miss_ are fractions of genes in or not in specific gene sets weighted by their correlations up to a given position in the rank; p values were estimated from random permutation tests by comparing random ES versus observed ES [82]. For qRT-PCR (***Fig.S2C***), RNA isolation and DNAseI digestion was performed with spleen, LN and BM cells stored in RNA later using RNAqueous-4PCR kit per manufacturer protocol (Ambion/Life Technologies). RNA integrity was evaluated on a RNA Nano chip run on a Bioanalyzer 2100 (Agilent Technologies); RNA integrity numbers (RIN) for all samples were 8.2-9.6. The cDNA first strand transcription was performed using 370ng of total RNA with the iScript cDNA synthesis kit following manufacturer protocol (BioRad). *Il7* and *Il15* primers were obtained from PrimerBank (http://pga.mgh.harvard.edu/primerbank/). Primer sequences for the SYBR green qPCR were as follows: *Il7* (PrimerBankID 6680433a1) forward (5-TTCCTCCACTGATCCTTGTTCT-3) & reverse (5-AGCAGCTTCC TTTGTATCATCAC-3), *Il15* (PrimerBankID 6680407a1) forward (5-ACATCCATCTCGTGCTACTTGT-3) & reverse (5-GCCTCTGTTTTAGGGAGACCT-3), *Gapdh* forward (5-AATGAAGGGGTCATTGATGG-3) & reverse (5-AAGGTGAAGGTCGGAGTCAA-3). Quantitative PCR was performed on a Roche LightCycler 480II Real Time PCR instrument, using PerfeCta SYBR Green (Quanta Biosciences). PCR was carried out in a 20ul volume and a final concentration of 1X reaction buffer, 385nM forward and reverse primers and 1.0ul cDNA reaction. Four log10 dilutions of pooled sample cDNA template were prepared and used for primer validation and standard curve reference. All sample reactions were performed in triplicate with NTC reactions for all primer sets on a single plate. PCR cycling parameters were as follows: hot-start at 95^o^C for 2min30sec, 45 cycles of 95^o^C for 15sec, 60^o^C for 35sec, followed by a dissociation curve measurement from 65^o^C to 95^o^C. Relative comparison analysis with efficiency correction was performed using the LC480II data collection software release 1.5.0.39 SP4. Melt curve analysis for all assays verified single product amplification and absence of primer dimers. NTC reactions for all primer sets were >5Cq from all control and unknown samples.

### In vitro *survival assays*

Single cell suspensions prepared from spleen or lympholyte-purified (Cedarlane) PBMCs were cultured for 12-48h in RPMI supplemented with 10% FCS but in the absence of added growth or survival factors (***Fig.2B/D***); in some cases, titrated amounts of pharmacological inhibitors ABT-737 (Abbott), C75 (Cayman), atglistatin (Cayman), chloroquine (Sigma) or vehicle were added to cultures (***Figs.2H, 4H/I & S3C-D***). CD8^+^T cell survival was subsequently determined by combined CD8α, congenic marker, MHC-I tetramer or CD44, and viability stains (Annexin V/propidium iodide [PI] or 7AAD, or Zombie dyes). Absolute numbers of viable CD8^+^T cell subsets were calculated using Countess (Invitrogen) or Vi-Cell (Beckmann Coulter) automated cell counters.

### In vivo *homing assays*

For competitive homing assays, splenic p14 T_M_ were enriched from young and old LCMV-immune p14 chimeras, differentially labeled with CFSE, mixed at a ratio of 1:1 and, depending on experiments, populations containing 1.1-4.2×10^5^ p14 T_M_ each were injected i.v. into B6 recipients; 42-48h later, transferred p14 T_M_ were retrieved and enumerated in LNs and other tissues (***Figs.6E & S5B***). Homing assays using endogenously generated CD8^+^T_M_ populations were conducted in an analogous fashion using 5-6×10^4^ D^b^NP_396_^+^ CD8^+^T_M_ each, B6.CD45.1 recipients and retrieval of donor cells from various tissues 20h later (***Fig.7E***). In some cases, mixed donor populations were incubated for 1h in complete RPMI (1.5×10^7^ cells/ml) in the presence or absence of 25ng/ml pertussis toxin (RnD Systems) prior to washes and transfer (***Fig.S5B***). For trafficking studies under conditions of CD62L blockade (***Fig.7A & S5C***), B6 mice were treated with a single i.p. injection of 200μg αCD62L (MEL-14) or rIgGa control (RTK2758) 2h before transfer of ∼5×10^5^ aged p14 T_M_ and retrieval 48h later. Further details about all antibodies are provided in ***Table S1***.

### Statistical analyses

Data handling, analysis and graphic representation was performed using Prism 6.0c (GraphPad Software). All data summarized in bar and line diagrams are expressed as mean ± 1 standard error (SEM), and asterisks indicate statistical differences calculated by Student’s t-test (unpaired or paired), or one-way ANOVA with Dunnett’s multiple comparisons test, and adopt the following convention: *: p<0.05, **: p<0.01 and ***: p<0.001.

## ACKNOWLEDGEMENTS

We wish to thank the NIH Tetramer Core Facility for provision of biotinylated MHC:peptide monomers, Dr.D. Hildeman for advice about the use of ABT-737, Dr. C. Buettner for provision of atglistatin, R. Wong for help with qRT-PCR analyses, and Dr. P. Marrack for the gift of the BIM antibody. This work was supported by NIH AG026518 and AI093637, JDRF CDA 2-2007-240, and DERC P30-DK057516 (DH), American Heart Association 13SDG14510023 (ETC), and NIH training grants T32 AI07405, T32 AI052066 and T32 DK007792 (BD); the funders had no role in study design, data collection and analysis, decision to publish, or preparation of the manuscript, and the authors declare no competing financial interests. Due to the wide-ranging nature of topics discussed herein we have frequently relied on the citation of review rather than original research articles and wish to apologize to the authors whose work is not explicitly mentioned.

## INDEX OF FIGURES, SUPPLEMENTARY FIGURES AND SUPPLEMENTARY TABLES

## I.FIGURES 1-7

1:Temporal regulation of major survival-associated components by aging CD8^+^T_M_.

2:Life & death of aging CD8^+^T_M_.

3:CD127/CD122 expression, signaling and homeostatic proliferation of aging CD8^+^T_M_

4:Metabolic adaptations of aging CD8^+^T_M_.

5:Increasing abundance and accelerated maturation of aging CD8^+^T_M_ in the splenic WP.

6:Progressive accumulation of aging CD8^+^T_M_ in peripheral LNs.

7:Redistribution of aging CD8^+^T_M_ from NLTs to lymphoid tissues.

## II.SUPPLEMENTARY FIGURES S1-S5

S1:Temporal regulation of survival- and apoptosis-related gene expression by p14 T_E/M_.

S2:Homeostasis of aging CD8^+^T_M_: staining controls, cell cycle GSEA, and cytokine mRNA levels. S3:Metabolic adaptations of aging CD8^+^T_M_.

S4:Phenotypic properties of RP and WP CD8^+^T_M_ populations in young and old mice.

S5:Chemokine receptor-dependent and CD62L-independent trafficking of young and old p14 T_M_.

## III.SUPPLEMENTARY TABLE S1

S1:Reagents & Materials

---

**Figure S1.**
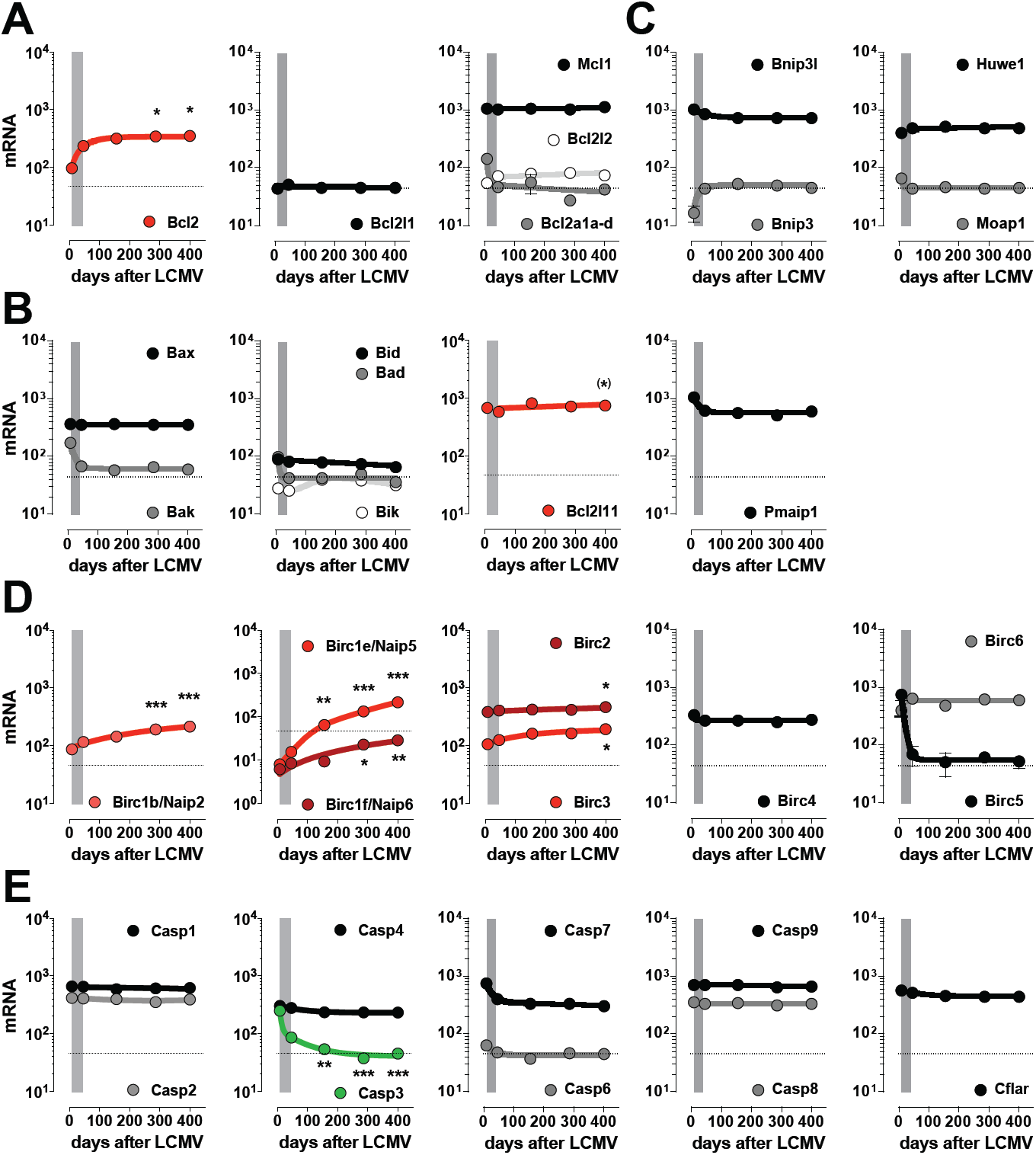
*Temporal regulation of survival- and apoptosis-related gene expression by p14 T*_*E/M*_. Transcriptional analyses were conducted with p14 T_E_ (day 8) and T_M_ (d46, d156, d286 and d400) purified from LCMV-challenged p14 chimeras and processed directly *ex vivo* for microarray hybridization as detailed in ref.^1^. The panels depict specific mRNA expression patterns of p14 T_E/M_ as a function of time after LCMV challenge, and the vertical gray bars indicate the transition period from T_E_ stage (d8) to early T_M_ stage (d42). **A.,** Bcl-2 family group IA (anti-apoptotic); **B.,** Bcl-2 family group IB (pro-apoptotic); **C.,** Bcl-2 family group IC (BH3-like contenders); **D.,** inhibitor of apoptosis proteins (IAPs, involved in the regulation of caspases, apoptosis, inflammatory signaling and immunity); **E.,** caspases. The data shown here for *Bcl2*, *Bcl2l1* (Bcl-x_L_), *Bcl2l11* (BIM) and *Casp3* are also displayed in ***Fig.1C/D***. All data are SEM with n≥3 individual mice/time point and asterisks indicate statistical significance comparing young (d40) and older (≥d156) p14 T_M_ using one-way ANOVA with Dunnett’s multiple comparisons test unless noted otherwise (*, p<0.05; **, p<0.01; ***, p<0.001; ^(^*^)^, significance of differential *Bcl2l11* expression comparing d46 and d400 by Student’s t-test but not ANOVA). For easier identification, significant differences emerging over the course of the memory phase are highlighted in red (up-regulation) or green (down-regulation).

**Figure S2.**
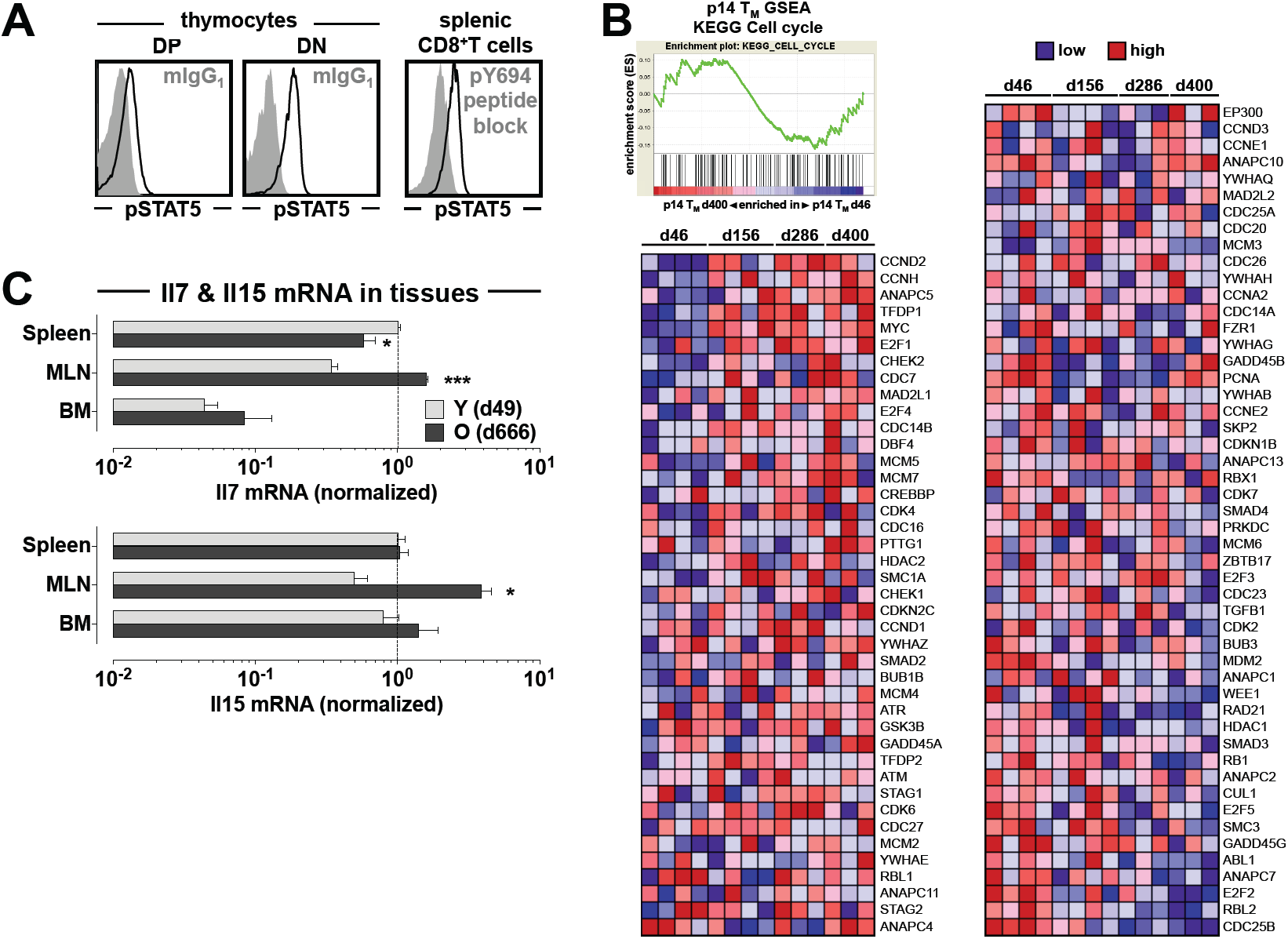
*Homeostasis of aging CD8*^*+*^*T*_*M*_*: staining controls, cell cycle GSEA, and cytokine mRNA levels.* **A.,** left: control stains (black tracing: pSTAT5, gray histograms: mIgG1 isotype) documenting differential constitutive pSTAT5 levels in DP *vs.* DN thymocytes in agreement with Van De Wiele *et al.*, J. Immunol. 172: 4235-4244, and thus absence of elevated non-specific staining using the pSTAT5 clone 47 antibody; right: *ex vivo* pSTAT5 stains of splenic CD8^+^T_M_ (d203); the blocking control (gray histogram) was performed by pre-incubation of the pSTAT5 antibody with an excess of pY694 peptide. **B.,** GSEAs were conducted for aging p14 T_M_ as detailed in Methods and demonstrate a non-significant negative enrichment for the cell cycle-associated KEGG gene set (NES = -0.68). **C.,** RNA was extracted from total spleen, MLN and BM cells obtained from young (d49) and old (d666) LCMV-immune p14 chimeras and analyzed by qRT-PCR as detailed in Methods; asterisks indicate significantly different *Il7* or *Il15* expression levels in respective young *vs.* old tissues (data from 1 of 2 similar experiments). All data are SEM with n≥3 individual mice.

**Figure S3.**
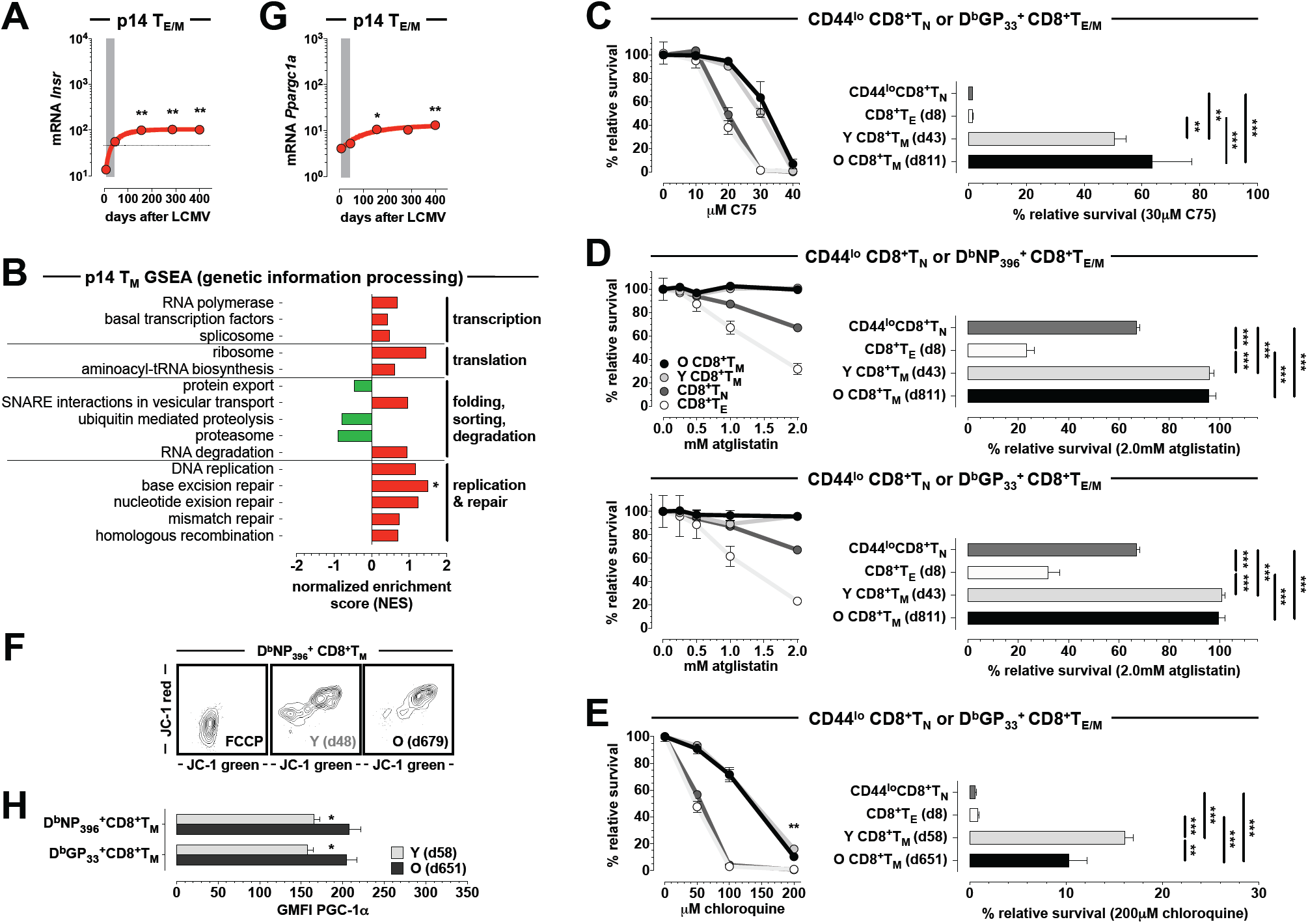
*Metabolic adaptations of aging CD8*^*+*^*T*_*M*_. **A.,** temporal regulation of *Insr* expression (p14 T_E/M_ microarray data). **B.,** summary of GSEAs that identify temporally regulated p14 T_M_-expressed gene sets within the KEGG category of “genetic information processing” (GIP; no other pathways in the GIP module demonstrated progressive temporal enrichment or depletion). **C.-E.,** CD8^+^T cell survival under conditions of lipogenesis or lipolysis inhibition. Spleen cells from naïve and indicated LCMV-infected B6 mice were cultured under conditions of “withdrawal apoptosis” in the presence of titrated amounts of indicated inhibitors or vehicle, and the survival of defined subsets (CD44^lo^CD8^+^T_N_ [dark gray], D^b^NP_396_ ^+^ and D^b^GP_33_ ^+^ CD8^+^T_E_ [white] as well as young [light gray] and aged [black] D^b^NP_396_ ^+^ and D^b^GP_33_ ^+^ CD8^+^T_M_) was quantified 24h later as detailed in Methods; given the differential survival of the different CD8^+^T cell populations in the absence of inhibitor, their relative survival under this condition was set for comparative purposes at 100%. All data are displayed as inhibitor titration curves (left) and under select conditions of inhibitor concentration (right). C., fatty acid synthase (FASN) inhibitor C75. D., adipose triglyceride lipase (ATGL) inhibitor atglistatin. E., inhibition of lysosomal acidifaction by chloroquine (n=4 mice/group; representative data from one of two experiments). **F.,** JC-1 stains of young and aged D^b^NP_396_^+^ CD8^+^T_M_. JC-1 is a membrane-permeant dye that exhibits potential-dependent accumulation in mitochondria as indicated by a green (∼529nm) to red (∼590) fluorescence shift; accordingly, mitochondrial depolarization decreases the red/green fluorescence intensity (incubation with FCCP prior to JC-1 stains was used as a control to dissipate the electrochemical proton gradient). **G.,** temporal regulation of *Ppargc1a* expression (p14 T_E/M_ microarray data). **H.,** PGC-1α (*Ppargc1a* gene product) expression by young and old LCMV-specific CD8^+^T_M_ (n=4 mice/group).

**Figure S4.**
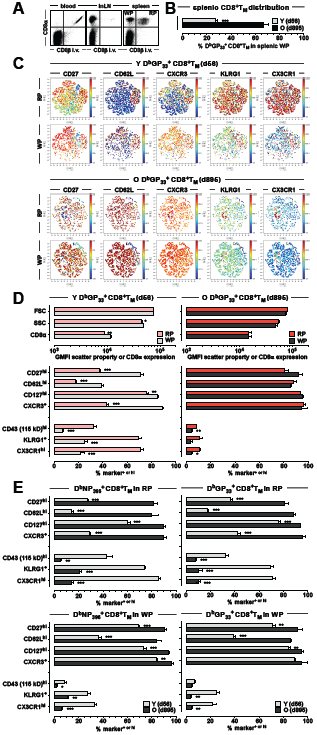
*Phenotypic properties of RP and WP CD8*^*+*^*T*_*M*_ *populations in young and old mice.* **A.,** intravascular CD8 staining labels blood-borne CD8^+^T cells, a very small fraction of LN cells (InLN: inguinal LN), and permits the distinction of splenic RP and WP cells (*cf.* refs.^55,56^). **B.,** relative fraction of young and old D^b^GP_33_^+^ CD8^+^T_M_ located in the splenic WP. **C.,** viSNE rendering of the D^b^GP_33_^+^ CD8^+^T_M_ phenotype space in RP *vs.* WP of young (top) and old (bottom) LCMV-immune mice. **D.,** individual phenotypic33 characteristics of D^b^GP_33_ ^+^ CD8^+^T_M_ RP and WP populations in young (left) and old (right) mice. **E.,** direct comparison of young and old D^b^NP_396_^+^ (left) and D^b^GP_33_ ^+^ (right) CD8^+^T_M_ in RP (top) and WP (bottom); n≥3 mice/time point.

**Figure S5.**
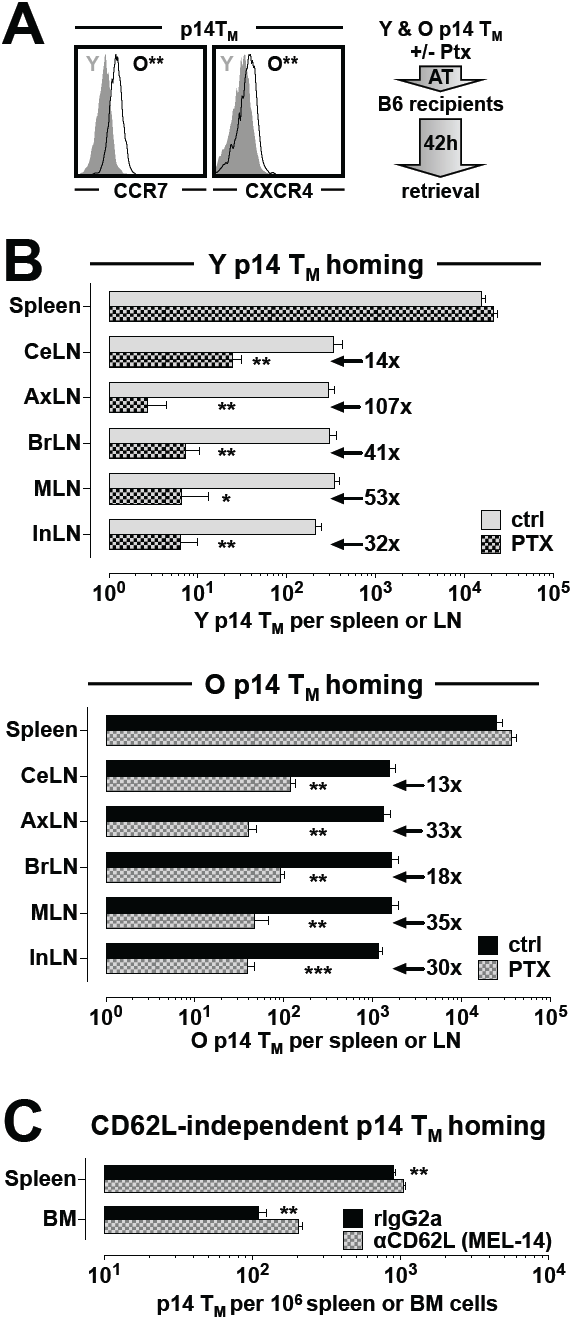
Chemokine receptor-dependent and CD62L-independent trafficking of young and old p14 TM. **A.,** left: histograms are gated on blood-borne p14 TM from young (d40, gray filled histograms) and old (d655, black tracings) LCMV-immune p14 chimeras; the asterisks indicate significant CCR7 and intracellular CXCR4 expression differences (n=3-5 mice). Right: experimental flow chart for p14 T_M_ trafficking experiments. Splenic p14 T_M_ populations obtained from young (d49) and old (d664) p14 chimeras were differentially labeled with CFSE, combined at a ratio of 1:1, and cultured for 1h at 37oC in the absence (control/ctrl) or presence of 25ng/ml pertussis toxin (Ptx) as described in Methods. Mixtures containing 4.2×10_5_ young and old p14 T_M_ each were subsequently transferred i.v. into B6 recipients and retrieved from various tissues 42h later. **B.,** enumeration of young (top) and old (bottom) p14 TM in spleen and lymph nodes; the values indicate the factor by which Ptx treatment reduced respective p14 T_M_ trafficking to indicated LNs (n=4 recipients each of mixed ctrl-or Ptx-treated p14 T_M_ populations). **C.,** aged p14 T_M_ (d630) were transferred into B6 recipients treated with rIgG2a isotype or ‹CD62L antibody, and retrieved 48h later as detailed in ***Fig.7A***. Note the enhanced accumulation of aged p14 T_M_ in spleen and BM under conditions of CD62L blockade, likely constituting a compensatory increase due to p14 T_M_ exclusion from LNs (n=4 mice/group).

**Table S1.**
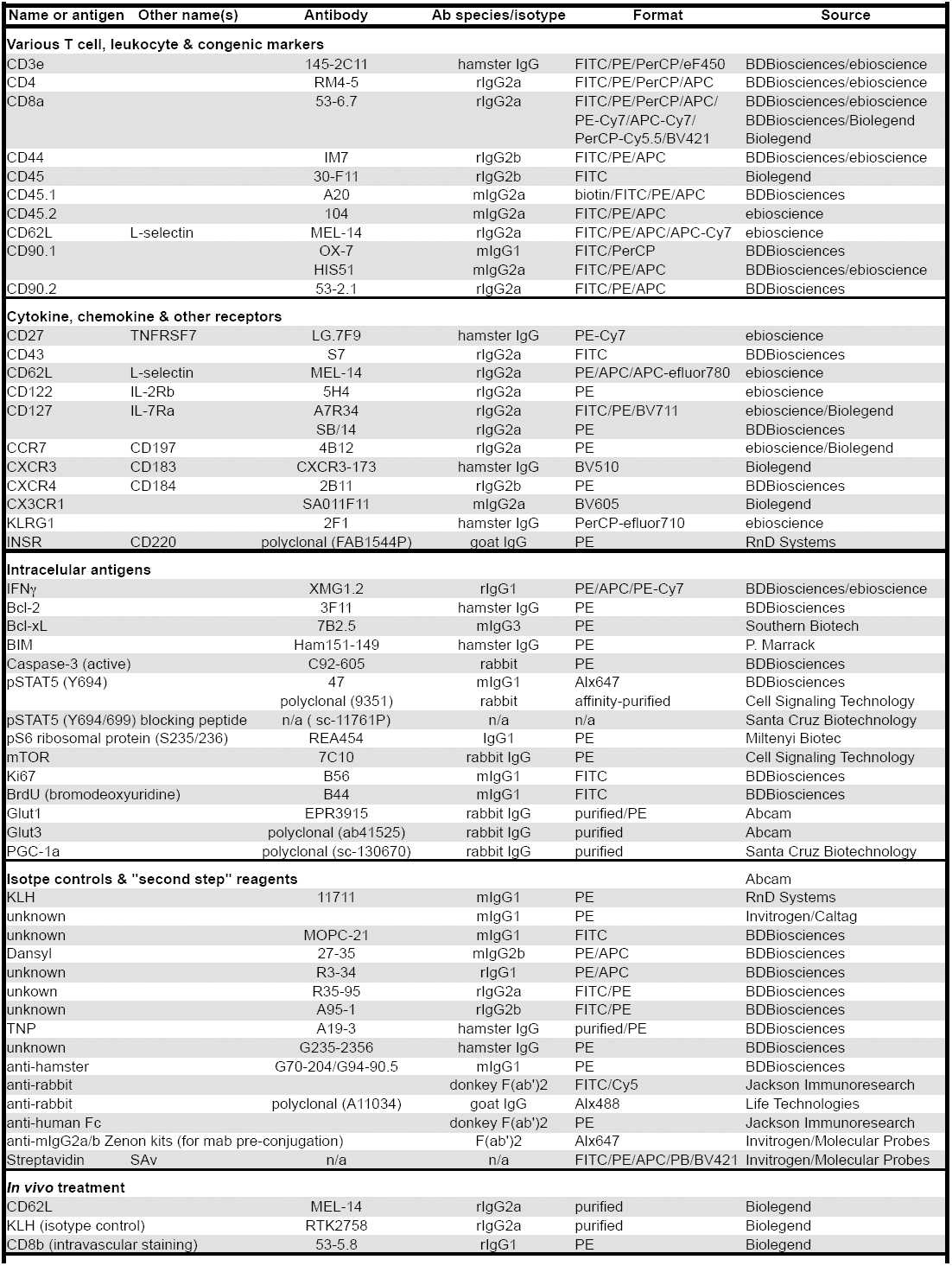
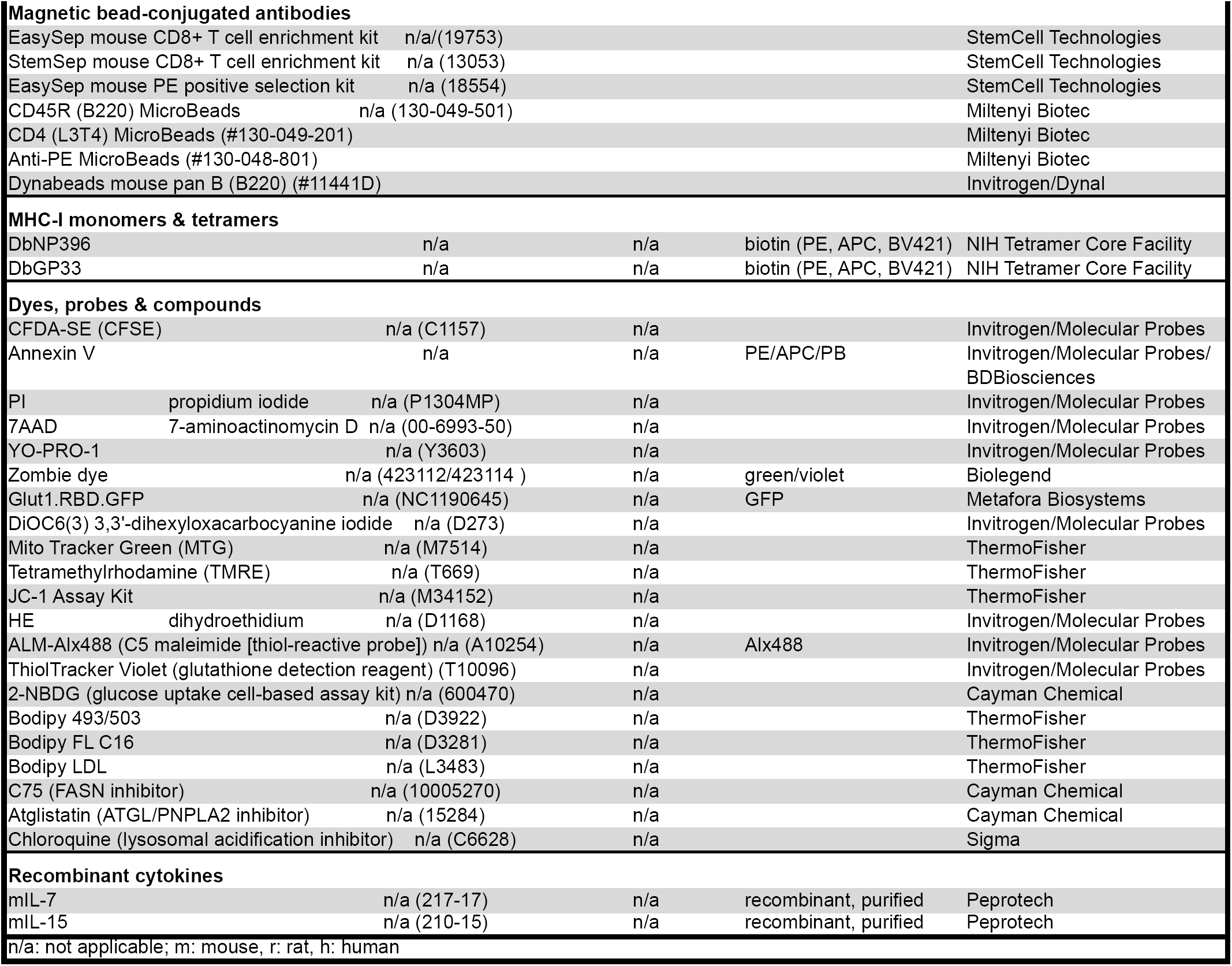
Reagents & Materials

## REFERENCES

1. Eberlein J, Davenport B, Nguyen TT, Victorino F, Karimpour-Fard A, Hunter LE, et al. Aging promotes acquisition of naïve-like CD8+ memory T cell traits and enhanced functionalities. J Clin Invest. 2016;106(10):3942–60. Epub September 12, 2016 doi:10.1172/JCI88546.

2. Roberts AD, Ely KH, Woodland DL. Differential contributions of central and effector memory T cells to recall responses. J Exp Med. 2005;202(1):123-33. PubMed PMID: 15983064.

3. Hikono H, Kohlmeier JE, Takamura S, Wittmer ST, Roberts AD, Woodland DL. Activation phenotype, rather than central-or effector-memory phenotype, predicts the recall efficacy of memory CD8+ T cells. J Exp Med. 2007;204(7):1625-36. PubMed PMID:: 17606632.

4. Martin MD, Condotta SA, Harty JT, Badovinac VP. Population dynamics of naive and memory CD8 T cell responses after antigen stimulations in vivo. J Immunol. 2012;188(3):1255–65. doi:10.4049/jimmunol.1101579. PubMed PMID:: 22205031; PubMed Central PMCID: PMC3262935.

5. Martin MD, Kim MT, Shan Q, Sompallae R, Xue HH, Harty JT, et al. Phenotypic and Functional Alterations in Circulating Memory CD8 T Cells with Time after Primary Infection. PLoS Pathog. 2015;11(10):e1005219. doi:10.1371/journal.ppat.1005219. PubMed PMID:: 26485703; PubMed Central PMCID: PMC4618693.

6. Akbar AN, Henson SM. Are senescence and exhaustion intertwined or unrelated processes that compromise immunity? Nat Rev Immunol. 2011;11(4):289–95. Epub 2011/03/26. doi:nri2959 [pii] 10.1038/nri2959. PubMed PMID:: 21436838.

7. Blackman MA, Woodland DL. The narrowing of the CD8 T cell repertoire in old age. Curr Opin Immunol. 2011;23(4):537–42. Epub 2011/06/10. doi:S0952-7915(11)00059-8 [pii] 10.1016/j.coi.2011.05.005. PubMed PMID:: 21652194; PubMed Central PMCID: PMC3163762.

8. Chou JP, Effros RB. T cell replicative senescence in human aging. Current pharmaceutical design. 2013;19(9):1680–98. PubMed PMID:: 23061726; PubMed Central PMCID: PMC3749774.

9. Nikolich-Zugich J. Aging of the T cell compartment in mice and humans: from no naive expectations to foggy memories. J Immunol. 2014;193(6):2622–9. doi:10.4049/jimmunol.1401174. PubMed PMID:: 25193936; PubMed Central PMCID: PMC4157314.

10. Goronzy JJ, Weyand CM. Successful and Maladaptive T Cell Aging. Immunity. 2017;46(3):364–78. doi:10.1016/j.immuni.2017.03.010. PubMed PMID:: 28329703.

11. Sallusto F, Lanzavecchia A, Araki K, Ahmed R. From vaccines to memory and back. Immunity. 2010;33(4):451–63. Epub 2010/10/30. doi:S1074-7613(10)00368-7 [pii] 10.1016/j.immuni.2010.10.008. PubMed PMID:: 21029957.

12. Walker JM, Slifka MK. Longevity of T-cell memory following acute viral infection. Adv Exp Med Biol. 2010;684:96-107. Epub 2010/08/28. PubMed PMID:: 20795543.

13. Murali-Krishna K, Lau LL, Sambhara S, Lemonnier F, Altman J, Ahmed R. Persistence of memory CD8 T cells in MHC class I-deficient mice. Science. 1999;286(5443):1377–81. PubMed PMID:: 10558996.

14. Homann D, Teyton L, Oldstone MB. Differential regulation of antiviral T-cell immunity results in stable CD8+ but declining CD4+ T-cell memory. Nat Med. 2001;7(8):913–9. PubMed PMID:: 11479623.

15. Surh CD, Sprent J. Homeostasis of naive and memory T cells. Immunity. 2008;29(6):848–62. Epub 2008/12/23. doi:S1074-7613(08)00506-2 [pii] 10.1016/j.immuni.2008.11.002. PubMed PMID:: 19100699.

16. Wojciechowski S, Tripathi P, Bourdeau T, Acero L, Grimes HL, Katz JD, et al. Bim/Bcl-2 balance is critical for maintaining naive and memory T cell homeostasis. J Exp Med. 2007;204(7):1665–75. PubMed PMID:: 17591857.

17. Wherry EJ, Teichgraber V, Becker TC, Masopust D, Kaech SM, Antia R, et al. Lineage relationship and protective immunity of memory CD8 T cell subsets. Nat Immunol. 2003;4(3):225–34. PubMed PMID:: 12563257.

18. Kurtulus S, Tripathi P, Moreno-Fernandez ME, Sholl A, Katz JD, Grimes HL, et al. Bcl-2 allows effector and memory CD8+ T cells to tolerate higher expression of Bim. J Immunol. 2011;186(10):5729–37. Epub 2011/04/01. doi:jimmunol.1100102 [pii] 4049/jimmunol.1100102. PubMed PMID:: 21451108.

19. Gyrd-Hansen M, Meier P. IAPs: from caspase inhibitors to modulators of NF-kappaB, inflammation and cancer. Nature reviews Cancer. 2010;10(8):561–74. doi:10.1038/nrc2889. PubMed PMID:: 20651737.

20. Gentle IE, Moelter I, Lechler N, Bambach S, Vucikuja S, Hacker G, et al. Inhibitor of apoptosis proteins (IAPs) are required for effective T cell expansion/survival during anti-viral immunity in mice. Blood. 2013. doi:10.1182/blood-2013-01-479543. PubMed PMID:: 24335231.

21. Grayson JM, Laniewski NG, Lanier JG, Ahmed R. Mitochondrial potential and reactive oxygen intermediates in antigen-specific CD8+ T cells during viral infection. J Immunol. 2003;170(9):4745–51. PubMed PMID:: 12707355.

22. Salminen A, Ojala J, Kaarniranta K. Apoptosis and aging: increased resistance to apoptosis enhances the aging process. Cell Mol Life Sci. 2011;68(6):1021–31. Epub 2010/12/01. doi:10.1007/s00018-010-0597-y. PubMed PMID:: 21116678.

23. Miller RA. Aging and immune function. In: Paul W, editor. Fundamental Immunology. 4th ed: Lippincott-Raven Publishers; 1999. p. 947–66.

24. Kim HJ, Nel AE. The role of phase II antioxidant enzymes in protecting memory T cells from spontaneous apoptosis in young and old mice. J Immunol. 2005;175(5):2948–59. Epub 2005/08/24. doi:175/5/2948 [pii]. PubMed PMID:: 16116181.

25. Peters T, Weiss JM, Sindrilaru A, Wang H, Oreshkova T, Wlaschek M, et al. Reactive oxygen intermediate-induced pathomechanisms contribute to immunosenescence, chronic inflammation and autoimmunity. Mech Ageing Dev. 2009;130(9):564–87. Epub 2009/07/28. doi:S0047-6374(09)00103-1 [pii] 10.1016/j.mad.2009.07.003. PubMed PMID:: 19632262.

26. Mak TW, Grusdat M, Duncan GS, Dostert C, Nonnenmacher Y, Cox M, et al. Glutathione Primes T Cell Metabolism for Inflammation. Immunity. 2017;46(4):675–89. doi:10.1016/j.immuni.2017.03.019. PubMed PMID:: 28423341.

27. Sahaf B, Heydari K, Herzenberg LA. Lymphocyte surface thiol levels. Proc Natl Acad Sci U S A. 2003;100(7):4001–5. PubMed PMID:: 12642656.

28. Oltersdorf T, Elmore SW, Shoemaker AR, Armstrong RC, Augeri DJ, Belli BA, et al. An inhibitor of Bcl-2 family proteins induces regression of solid tumours. Nature. 2005;435(7042):677–81. PubMed PMID:: 15902208.

29. Masopust D, Ha SJ, Vezys V, Ahmed R. Stimulation history dictates memory CD8 T cell phenotype: implications for prime-boost vaccination. J Immunol. 2006;177(2):831–9. PubMed PMID:: 16818737.

30. Song J, So T, Cheng M, Tang X, Croft M. Sustained survivin expression from OX40 costimulatory signals drives T cell clonal expansion. Immunity. 2005;22(5):621–31. Epub 2005/05/17. doi:S1074-7613(05)00107-X [pii] 10.1016/j.immuni.2005.03.012. PubMed PMID:: 15894279.

31. Dunkle A, Dzhagalov I, Gordy C, He YW. Transfer of CD8+ T cell memory using Bcl-2 as a marker. J Immunol. 2013;190(3):940–7. doi:10.4049/jimmunol.1103481. PubMed PMID:: 23269245.

32. Choo DK, Murali-Krishna K, Anita R, Ahmed R. Homeostatic turnover of virus-specific memory CD8 T cells occurs stochastically and is independent of CD4 T cell help. J Immunol. 2010;185(6):3436–44. doi:10.4049/jimmunol.1001421. PubMed PMID:: 20733203.

33. Kemp RA, Pearson CF, Cornish GH, Seddon BP. Evidence of STAT5-dependent and-independent routes to CD8 memory formation and a preferential role for IL-7 over IL-15 in STAT5 activation. Immunol Cell Biol. 2010;88(2):213–9. Epub 2009/12/02. doi:icb200995 [pii] 10.1038/icb.2009.95. PubMed PMID:: 19949423; PubMed Central PMCID: PMC2842934.

34. Pandey A, Ozaki K, Baumann H, Levin SD, Puel A, Farr AG, et al. Cloning of a receptor subunit required for signaling by thymic stromal lymphopoietin. Nat Immunol. 2000;1(1):59–64. PubMed PMID:: 10881176.

35. Willinger T, Freeman T, Hasegawa H, McMichael AJ, Callan MF. Molecular signatures distinguish human central memory from effector memory CD8 T cell subsets. J Immunol. 2005;175(9):5895–903. Epub 2005/10/21. doi:175/9/5895 [pii]. PubMed PMID:: 16237082.

36. Hand TW, Cui W, Jung YW, Sefik E, Joshi NS, Chandele A, et al. Differential effects of STAT5 and PI3K/AKT signaling on effector and memory CD8 T-cell survival. Proc Natl Acad Sci U S A. 2010;107(38):16601–6. Epub 2010/09/09. doi:1003457107 [pii] 10.1073/pnas.1003457107. PubMed PMID:: 20823247; PubMed Central PMCID: PMC2944719.

37. Zhang X, Fujii H, Kishimoto H, LeRoy E, Surh CD, Sprent J. Aging leads to disturbed homeostasis of memory phenotype CD8(+) cells. J Exp Med. 2002;195(3):283–93. Epub 2002/02/06. PubMed PMID:: 11828003; PubMed Central PMCID: PMC2193587.

38. Lenz DC, Kurz SK, Lemmens E, Schoenberger SP, Sprent J, Oldstone MB, et al. IL-7 regulates basal homeostatic proliferation of antiviral CD4+T cell memory. Proc Natl Acad Sci U S A. 2004;101(25):9357–62. PubMed PMID:: 15197277.

39. Marzo AL, Klonowski KD, Le Bon A, Borrow P, Tough DF, Lefrancois L. Initial T cell frequency dictates memory CD8+ T cell lineage commitment. Nat Immunol. 2005;6(8):793–9. PubMed PMID:: 16025119.

40. Buck MD, O’Sullivan D, Pearce EL. T cell metabolism drives immunity. J Exp Med. 2015;212(9):1345–60. doi:10.1084/jem.20151159. PubMed PMID:: 26261266; PubMed Central PMCID: PMCPMC4548052.

41. Chapman NM, Chi H. mTOR Links Environmental Signals to T Cell Fate Decisions. Frontiers in immunology. 2014;5:686. doi:10.3389/fimmu.2014.00686. PubMed PMID:: 25653651; PubMed Central PMCID: PMCPMC4299512.

42. Pollizzi KN, Powell JD. Regulation of T cells by mTOR: the known knowns and the known unknowns. Trends Immunol. 2015;36(1):13–20. doi:10.1016/j.it.2014.11.005. PubMed PMID:: 25522665; PubMed Central PMCID: PMCPMC4290883.

43. Phan AT, Doedens AL, Palazon A, Tyrakis PA, Cheung KP, Johnson RS, et al. Constitutive Glycolytic Metabolism Supports CD8+ T Cell Effector Memory Differentiation during Viral Infection. Immunity. 2016;45(5):1024–37. doi:10.1016/j.immuni.2016.10.017. PubMed PMID:: 27836431; PubMed Central PMCID: PMCPMC5130099.

44. Bengsch B, Johnson AL, Kurachi M, Odorizzi PM, Pauken KE, Attanasio J, et al. Bioenergetic Insufficiencies Due to Metabolic Alterations Regulated by the Inhibitory Receptor PD-1 Are an Early Driver of CD8(+) T Cell Exhaustion. Immunity. 2016;45(2):358–73. doi:10.1016/j.immuni.2016.07.008. PubMed PMID:: 27496729; PubMed Central PMCID: PMCPMC4988919.

45. O’Sullivan D, van der Windt GJ, Huang SC, Curtis JD, Chang CH, Buck MD, et al. Memory CD8(+) T cells use cell-intrinsic lipolysis to support the metabolic programming necessary for development. Immunity. 2014;41(1):75–88. doi:10.1016/j.immuni.2014.06.005. PubMed PMID:: 25001241; PubMed Central PMCID: PMCPMC4120664.

46. Macintyre AN, Gerriets VA, Nichols AG, Michalek RD, Rudolph MC, Deoliveira D, et al. The glucose transporter Glut1 is selectively essential for CD4 T cell activation and effector function. Cell Metab. 2014;20(1):61–72. doi:10.1016/j.cmet.2014.05.004. PubMed PMID:: 24930970; PubMed Central PMCID: PMCPMC4079750.

47. Proud CG. Regulation of protein synthesis by insulin. Biochemical Society transactions. 2006;34(Pt 2):213–6. doi:10.1042/BST20060213. PubMed PMID:: 16545079.

48. Lochner M, Berod L, Sparwasser T. Fatty acid metabolism in the regulation of T cell function. Trends Immunol. 2015;36(2):81–91. doi:10.1016/j.it.2014.12.005. PubMed PMID:: 25592731.

49. Masopust D, Lefrancois L. CD8 T-cell memory: the other half of the story. Microbes Infect. 2003;5(3):221–6. PubMed PMID:: 12681411.

50. Klonowski KD, Williams KJ, Marzo AL, Blair DA, Lingenheld EG, Lefrancois L. Dynamics of blood-borne CD8 memory T cell migration in vivo. Immunity. 2004;20(5):551–62. PubMed PMID:: 15142524.

51. Sarkar S, Teichgraber V, Kalia V, Polley A, Masopust D, Harrington LE, et al. Strength of stimulus and clonal competition impact the rate of memory CD8 T cell differentiation. J Immunol. 2007;179(10):6704–14. Epub 2007/11/06. doi:179/10/6704 [pii]. PubMed PMID:: 17982060.

52. Kohlmeier JE, Woodland DL. Immunity to respiratory viruses. Annu Rev Immunol. 2009;27:61-82. Epub 2008/10/29. doi:10.1146/annurev.immunol.021908.132625. PubMed PMID:: 18954284.

53. Nolz JC, Starbeck-Miller GR, Harty JT. Naive, effector and memory CD8 T-cell trafficking: parallels and distinctions. Immunotherapy. 2011;3(10):1223–33. doi:10.2217/imt.11.100. PubMed PMID:: 21995573; PubMed Central PMCID: PMC3214994.

54. Jung YW, Rutishauser RL, Joshi NS, Haberman AM, Kaech SM. Differential localization of effector and memory CD8 T cell subsets in lymphoid organs during acute viral infection. J Immunol. 2010;185(9):5315–25. doi:10.4049/jimmunol.1001948. PubMed PMID:: 20921525.

55. Anderson KG, Mayer-Barber K, Sung H, Beura L, James BR, Taylor JJ, et al. Intravascular staining for discrimination of vascular and tissue leukocytes. Nat Protoc. 2014;9(1):209–22. doi:10.1038/nprot.2014.005. PubMed PMID:: 24385150; PubMed Central PMCID: PMC4428344.

56. Steinert EM, Schenkel JM, Fraser KA, Beura LK, Manlove LS, Igyarto BZ, et al. Quantifying Memory CD8 T Cells Reveals Regionalization of Immunosurveillance. Cell. 2015;161(4):737–49. doi:10.1016/j.cell.2015.03.031. PubMed PMID:: 25957682; PubMed Central PMCID: PMC4426972.

57. Aw D, Hilliard L, Nishikawa Y, Cadman ET, Lawrence RA, Palmer DB. Disorganization of the splenic microanatomy in ageing mice. Immunology. 2016;148(1):92–101. doi:10.1111/imm.12590. PubMed PMID:: 26840375; PubMed Central PMCID: PMCPMC4819137.

58. Scimone ML, Felbinger TW, Mazo IB, Stein JV, Von Andrian UH, Weninger W. CXCL12 mediates CCR7-independent homing of central memory cells, but not naive T cells, in peripheral lymph nodes. J Exp Med. 2004;199(8):1113–20. PubMed PMID:: 15096537.

59. von Andrian UH, Mempel TR. Homing and cellular traffic in lymph nodes. Nat Rev Immunol. 2003;3(11):867–78. Epub 2003/12/12. doi:10.1038/nri1222nri1222 [pii]. PubMed PMID:: 14668803.

60. Nolz JC, Harty JT. Protective capacity of memory CD8+ T cells is dictated by antigen exposure history and nature of the infection. Immunity. 2011;34(5):781–93. Epub 2011/05/10. doi:S1074-7613(11)00174-9 [pii] 10.1016/j.immuni.2011.03.020. PubMed PMID:: 21549619; PubMed Central PMCID: PMC3103642.

61. Van den Broeck W, Derore A, Simoens P. Anatomy and nomenclature of murine lymph nodes: Descriptive study and nomenclatory standardization in BALB/cAnNCrl mice. J Immunol Methods. 2006;312(1-2):12-9. Epub 2006/04/21. doi:S0022-1759(06)00064-0 [pii] 10.1016/j.jim.2006.01.022. PubMed PMID:: 16624319.

62. Mueller SN, Gebhardt T, Carbone FR, Heath WR. Memory T cell subsets, migration patterns, and tissue residence. Annu Rev Immunol. 2013;31:137-61. doi:10.1146/annurev-immunol-032712-095954. PubMed PMID:: 23215646.

63. Jiang X, Clark RA, Liu L, Wagers AJ, Fuhlbrigge RC, Kupper TS. Skin infection generates non-migratory memory CD8+ T(RM) cells providing global skin immunity. Nature. 2012;483(7388):227–31. doi:10.1038/nature10851. PubMed PMID:: 22388819; PubMed Central PMCID: PMC3437663.

64. Schenkel JM, Masopust D. Tissue-resident memory T cells. Immunity. 2014;41(6):886–97. doi:10.1016/j.immuni.2014.12.007. PubMed PMID:: 25526304; PubMed Central PMCID: PMC4276131.

65. Xu RH, Fang M, Klein-Szanto A, Sigal LJ. Memory CD8+ T cells are gatekeepers of the lymph node draining the site of viral infection. Proc Natl Acad Sci U S A. 2007;104(26):10992–7. PubMed PMID:: 17578922.

66. Gasteiger G, Ataide M, Kastenmuller W. Lymph node-an organ for T-cell activation and pathogen defense. Immunol Rev. 2016;271(1):200–20. doi:10.1111/imr.12399. PubMed PMID:: 27088916.

67. Chaix J, Nish SA, Lin WH, Rothman NJ, Ding L, Wherry EJ, et al. Cutting edge: CXCR4 is critical for CD8+ memory T cell homeostatic self-renewal but not rechallenge self-renewal. J Immunol. 2014;193(3):1013–6. doi:10.4049/jimmunol.1400488. PubMed PMID:: 24973450; PubMed Central PMCID: PMC4108510.

68. Di Rosa F, Gebhardt T. Bone Marrow T Cells and the Integrated Functions of Recirculating and Tissue-Resident Memory T Cells. Frontiers in immunology. 2016;7:51. doi:10.3389/fimmu.2016.00051. PubMed PMID:: 26909081; PubMed Central PMCID: PMC4754413.

69. Kimura MY, Pobezinsky LA, Guinter TI, Thomas J, Adams A, Park JH, et al. IL-7 signaling must be intermittent, not continuous, during CD8(+) T cell homeostasis to promote cell survival instead of cell death. Nat Immunol. 2013;14(2):143–51. doi:10.1038/ni.2494. PubMed PMID:: 23242416; PubMed Central PMCID: PMC3552087.

70. Westera L, van Hoeven V, Drylewicz J, Spierenburg G, van Velzen JF, de Boer RJ, et al. Lymphocyte maintenance during healthy aging requires no substantial alterations in cellular turnover. Aging cell. 2015. doi:10.1111/acel.12311. PubMed PMID:: 25627171.

71. Marzo AL, Yagita H, Lefrancois L. Cutting edge: migration to nonlymphoid tissues results in functional conversion of central to effector memory CD8 T cells. J Immunol. 2007;179(1):36–40. PubMed PMID:: 17579018; PubMed Central PMCID: PMC2861291.

72. Kohlmeier JE, Miller SC, Woodland DL. Cutting edge: Antigen is not required for the activation and maintenance of virus-specific memory CD8+ T cells in the lung airways. J Immunol. 2007;178(8):4721–5. PubMed PMID:: 17404250.

73. Thome JJC, Yudanin N, Ohmura Y, Kubota M, Grinshpun B, Sathaliyawala T, et al. Spatial Map of Human T Cell Compartmentalization and Maintenance over Decades of Life. Cell. 2014;159:814–28.

74. Best JA, Blair DA, Knell J, Yang E, Mayya V, Doedens A, et al. Transcriptional insights into the CD8(+) T cell response to infection and memory T cell formation. Nat Immunol. 2013;14(4):404–12. doi:10.1038/ni.2536. PubMed PMID:: 23396170; PubMed Central PMCID: PMC3689652.

75. Doedens AL, Phan AT, Stradner MH, Fujimoto JK, Nguyen JV, Yang E, et al. Hypoxia-inducible factors enhance the effector responses of CD8(+) T cells to persistent antigen. Nat Immunol. 2013;14(11):1173–82. doi:10.1038/ni.2714. PubMed PMID:: 24076634; PubMed Central PMCID: PMC3977965.

76. Dominguez CX, Amezquita RA, Guan T, Marshall HD, Joshi NS, Kleinstein SH, et al. The transcription factors ZEB2 and T-bet cooperate to program cytotoxic T cell terminal differentiation in response to LCMV viral infection. J Exp Med. 2015;212(12):2041–56. doi:10.1084/jem.20150186. PubMed PMID:: 26503446; PubMed Central PMCID: PMC4647261.

77. Gray SM, Kaech SM, Staron MM. The interface between transcriptional and epigenetic control of effector and memory CD8(+) T-cell differentiation. Immunol Rev. 2014;261(1):157–68. doi:10.1111/imr.12205. PubMed PMID:: 25123283.

78. Gelderman KA, Hultqvist M, Holmberg J, Olofsson P, Holmdahl R. T cell surface redox levels determine T cell reactivity and arthritis susceptibility. Proc Natl Acad Sci U S A. 2006;103(34):12831–6. PubMed PMID:: 16908843.

79. Hildeman DA, Mitchell T, Teague TK, Henson P, Day BJ, Kappler J, et al. Reactive oxygen species regulate activation-induced T cell apoptosis. Immunity. 1999;10(6):735–44. PubMed PMID:: 10403648.

80. Eberlein J, Nguyen TT, Victorino F, Golden-Mason L, Rosen HR, Homann D. Comprehensive assessment of chemokine expression profiles by flow cytometry. J Clin Invest. 2010;120(3):907–23. Epub 2010/03/04. doi:40645 [pii] 10.1172/JCI40645. PubMed PMID:: 20197626; PubMed Central PMCID: PMC2827956.

81. Amir el AD, Davis KL, Tadmor MD, Simonds EF, Levine JH, Bendall SC, et al. viSNE enables visualization of high dimensional single-cell data and reveals phenotypic heterogeneity of leukemia. Nat Biotechnol. 2013;31(6):545–52. doi:10.1038/nbt.2594. PubMed PMID:: 23685480; PubMed Central PMCID: PMC4076922.

82. Subramanian A, Tamayo P, Mootha VK, Mukherjee S, Ebert BL, Gillette MA, et al. Gene set enrichment analysis: a knowledge-based approach for interpreting genome-wide expression profiles. Proc Natl Acad Sci U S A. 2005;102(43):15545–50. Epub 2005/10/04. doi:0506580102 [pii] 1073/pnas.0506580102. PubMed PMID:: 16199517; PubMed Central PMCID: PMC1239896.

